# Multiple human enhancer RNAs contain long translated open reading frames

**DOI:** 10.1101/2025.04.18.649583

**Authors:** Pavel A. Vlasov, Koichi Ogami, Elizabeth Valenzuela, Risa Karakida Kawaguchi, Marko Jovanovic, James L. Manley

## Abstract

Enhancer RNAs (eRNAs) are transcribed by RNA polymerase II during enhancer activation but are typically rapidly degraded in the nucleus. During states of reduced RNA surveillance, however, eRNAs and other similar “noncoding” RNAs, including for example upstream antisense RNAs, are stabilized, and some are exported to the cytoplasm and can even be found on polysomes. Here, we report unexpectedly that ∼12% of human intergenic eRNAs contain long open reading frames (>300 nts), many of which can be actively translated, as determined by ribosome profiling, and produce proteins that accumulate in cells, as shown by mass spectrometry (MS) data. Focusing on the largest of the encoded proteins, which we designate as eORFs, and which can be up to ∼45 KDa, we found remarkably that most are highly basic, with pIs >11.5. This unusual chemistry reflects a striking overabundance of arginine residues, and occurs despite a relative paucity of lysines. Exogenous expression of the ten largest eORFs revealed that they accumulate stably in cells as full-length proteins, and most localize to the nucleus and associate with chromatin. Identification of interacting proteins by MS suggested possible roles for these proteins in several nuclear processes. The eORFs studied are well-conserved among primates, although they are largely absent from other mammals. Notably, several contain human-specific C-terminal extensions and display properties suggestive of de novo gene birth. In summary, we have discovered that a fraction of human eRNAs can function as mRNAs, revealing a new and unexpected role for these transcripts.

## Introduction

Enhancers are distal regulatory sequences that play important roles in expression of many genes across the human genome. These sites are typically marked by unique epigenetic signatures, such as histone H3 lysine 4 monomethylation and lysine 27 acetylation, that distinguish them from promoters and other regulatory sequences. Enhancers can be located within or outside of gene sequences and their activity varies greatly between different cell types, tissue types and development stages. Part of the process of enhancer activation and interaction with their target genes involves binding of RNA polymerase II (RNAP II) to the enhancer site (Kim and Shiekhattar 2015; Field and Adelman 2020; Barral and Dejardin 2023).

RNAP II transcribes a wide variety of RNAs, including both mRNAs and long noncoding (lnc) RNAs. At enhancer sites, RNAP II transcribes a class of lncRNAs called enhancer RNAs (eRNAs), which are typically relatively short and are often transcribed bidirectionally from active enhancers (De Santa et al. 2010; Kim and Shiekhattar 2015). While the functions of eRNAs are not fully understood, they have been found to play roles in gene activation through enhancer loop formation, chromatin remodeling, interacting with and recruiting transcription factors, and possibly mediating heterotypic interactions leading to phase separation during promoter activation (Hsieh et al. 2014; Lai and Shiekhattar 2014; Arnold et al. 2019; Sartorelli and Lauberth 2020). Additionally, eRNAs have been implicated in disease, for example eRNA-associated SNPs have been associated with Alzheimer’s disease (Kikuchi et al. 2019). Furthermore, not only are enhancers able to produce RNAs similar to mRNAs, but some mRNA promoters have been found to exhibit enhancer-like function (Medina-Rivera et al. 2018). And conversely, transcription of eRNAs from enhancers shares many properties with transcription from mRNA promoters, including promoter architecture and transcriptional pausing and release (Core et al. 2014; Henriques et al. 2018).

eRNAs are naturally highly unstable. These and other unstable lncRNAs are typically targeted by the Mtr4 RNA helicase for degradation by the nuclear exosome complex (Zinder and Lima 2017; Garland and Jensen 2020). These lncRNAs include for example upstream antisense RNAs (uaRNAs) and prematurely terminated RNAs (ptRNAs), which are transcribed at promoter sites by RNAP II. These lncRNA species share similarities with mRNAs, e.g., they all undergo 5’ capping, and some are also spliced and 3’ polyadenylated. Mtr4 is involved in multiple protein complexes that target different lncRNAs for degradation, such as the NEXT and TRAMP complexes (Lubas et al. 2011; Laffleur et al. 2017; Ogami et al. 2018; Lykke-Andersen et al. 2021; Gockert et al. 2022; Puno and Lima 2022; Rouviere et al. 2022). Depletion of Mtr4 or related quality control factors leads to accumulation of these transcripts, which allows them to be exported from the nucleus and in some cases translated (Fan et al. 2017; Ogami and Manley 2017; Ogami et al. 2017; Silla et al. 2018; Ogami and Suzuki 2021).

Translation of ORFs located outside of canonical mRNAs has been detected across many RNA species. This includes lncRNAs with already known functions, with most studies so far focused on small ORFs shorter than 300 nts (Ingolia et al. 2014; Erhard et al. 2018; Olexiouk et al. 2018). For example, a short ORF encoding a peptide expressed during neuronal development has been identified in the well-studied lncRNA *MALAT1* (Xiao et al. 2024). Some ORF-containing lncRNAs were found to be differentially expressed during cancer development across multiple unrelated cancer types such as colon and breast cancer, and their products include both tumor suppressing and oncogenic peptides (Zhou et al. 2021; Della Bella et al. 2022; Wen et al. 2024). For example, a group of lncRNAs was found to be dysregulated in certain cancers, and some translated to produce small antigenic peptides presented on MHC class I protein complexes, acting as antigens for an anti-tumor immune response (Barczak et al. 2023). Notably though, lncRNA translation is not limited to small ORFs, and can occur on RNAs with their own regulatory functions. For example, the nucleolar lncRNA *PAPAS* is transcribed antisense to rRNAs and is involved in modulating rRNA expression through multiple repression mechanisms (Zhao et al. 2016; Zhao et al. 2018). However, in humans and other primates, the transcript also encodes a 229 aa protein, RIEP, involved in nucleolar and mitochondrial heat shock stress responses (Feng et al. 2023).

In this study, we investigated the potential translatability of eRNAs globally in human cells. We first identified ORF-containing eRNA transcripts computationally, and found a surprisingly large number with ORFs >300 nts. Ribosome profiling following Mtr4 depletion was used to identify actively translated eRNAs, and those with the largest ORFs were analyzed further. Remarkably, the encoded proteins were almost all extremely basic, highly enriched in arginine residues, and predicted to exhibit mostly nuclear localization. Analysis of mass spectrometry (MS) databases revealed that some of these proteins accumulate endogenously in cells even in the absence of Mtr4 depletion, while exogenous expression confirmed their stability and primarily nuclear localization. Immunoprecipitation followed by MS provided evidence for association of the proteins with DNA or RNA-interacting protein complexes. Finally, these large ORFs are conserved in primates, although notably several have human-specific C-terminal extensions. Our results not only provide unexpected new insights into functions of eRNAs, but also, and more broadly, suggest the existence of a large reservoir of previously unknown protein-coding sequences that expand the human proteome in ways yet to be established.

## Results

### Identification of potentially translatable eRNAs

Our previous work and that of others has provided evidence that certain lncRNAs considered to be noncoding do in fact have the potential to be translated. In this study, we wished to examine globally how widespread this potential is among one such class of lncRNAs, eRNAs. To this end, we obtained candidate enhancer sites from the ENCODE Project Consortium (ENCODE Project Consortium 2012) and chose those where poly(A^+^) RNA-seq results indicated bidirectional transcription originating from them (Djebali et al. 2012). To avoid potentially misidentifying mRNAs as eRNAs, we selected only intergenic enhancer sites located more than 2.5 kb away from the nearest gene for further analysis. Based on these criteria, 20,946 putative enhancer sites producing 41,892 possible eRNA transcripts (two per site) were identified. A preliminary test for the presence of ORFs >300 nt was performed using orfipy (Singh and Wurtele 2021), with 5161 eRNAs (∼12%) containing at least one long ORF starting with ATG, or 9259 eRNAs (∼22%) when the alternative possible start codon CTG (Kearse and Wilusz 2017) was included. Predicted eRNA sequences were further examined for translation potential using the Coding Potential Assessment Tool, CPAT (Wang et al. 2013; Li and Wang 2021). Out of all eRNAs tested, 753 contained ORFs rated as having a coding potential of 50% or higher, ∼15% of all ATG ORF-containing eRNAs or 1.8% of all eRNAs (although see below).

### Ribosome profiling of Mtr4-depleted cells provides evidence for eRNA translation

We next investigated whether any of these eRNAs could actually be translated. To increase the likelihood of detecting these typically unstable, nuclear transcripts undergoing translation in the cytoplasm, we took advantage of our previous findings that knockdown (KD) of Mtr4 resulted in the stabilization, transport to the cytoplasm and translation of certain ua- and pt-RNAs (Ogami et al. 2017). We therefore performed ribosome profiling (Ingolia et al. 2009; Ingolia 2016) of Mtr4-depleted and control HEK293T cells to identify actively translated eRNAs (Figure 1A; see Methods for details). This analysis identified 923 total transcripts upregulated by at least 2-fold after Mtr4 KD, including 182 transcripts identified as eRNAs (Figure 1B). Other upregulated transcripts included both mRNAs, other lncRNAs such as uaRNAs, and novel transcripts predicted by StringTie (Supp. Fig. 1A). While published RNAs included both up- and down-regulated transcripts, all eRNAs that exhibited significant changes were upregulated (Supp. Fig. 1B).

**Figure 1:**
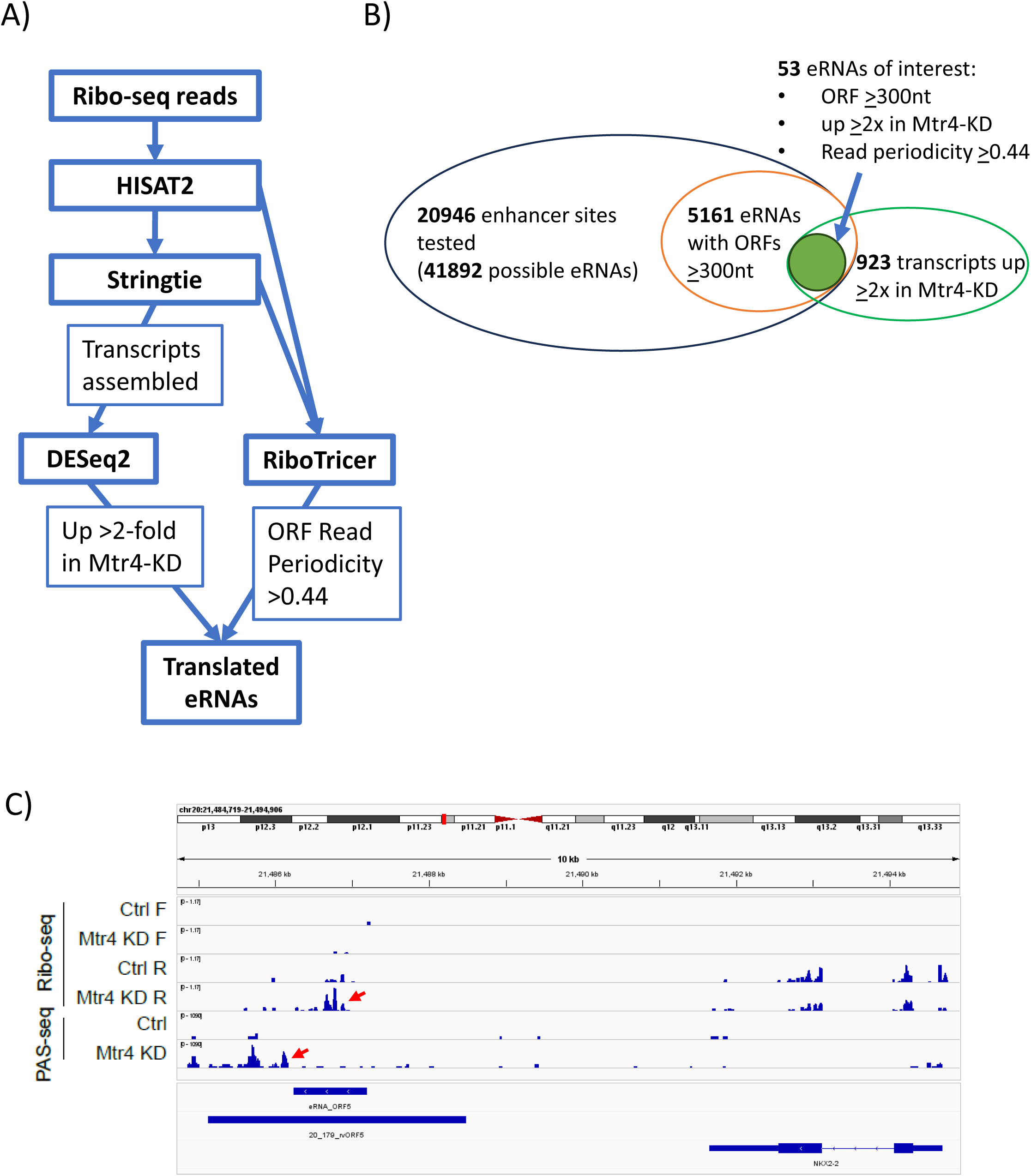
Identification of translated eRNAs by ribosome profiling. **A** Ribosome profiling data analysis pipeline. Reads were aligned by HISAT2. Transcripts were assembled using StringTie guided by the human transcriptome database and predicted eRNA sequences. Reads were quantified and compared between Mtr4-KD and control samples by DESeq2. eRNA ORFs were identified and their read periodicity was calculated by RiboTricer. **B** Summary of transcripts tested by StringTie and DeSEQ2 analysis of ribosome profiling reads. eRNA sequences predicted from 20946 predicted enhancer sites were tested alongside all published human transcriptome sequences. eRNAs were selected for further analysis based on the following criteria: upregulation by >2x in Mtr4-KD samples (from DeSEQ2), ORF size of >300 nt (from orfipy, CPAT, and RiboTricer), and read periodicity (from RiboTricer). 53 eRNAs met these critera and the 10 containing the largest ORFs were selected (ORFs 1-10). **C** IGV plot of ribosome profiling (reads from samples combined and sorted by direction) and PAS-seq reads at the site of ORF5, with both the enhancer site and ORF location shown below.

We next set out to obtain a high confidence list of translated eRNAs by examining properties of the polysome-associated eRNAs identified above. Among the eRNA transcripts with ORFs >300 nt, 53 passed the minimum read periodicity (phase score >0.44) calculated by Ribotricer for human ORF sequences (Choudhary et al. 2020). Read numbers and phase scores for all these eRNA ORFs, which we refer to as eORFs, were within the range observed for typical lower-expression level mRNAs, e.g. those encoding various ZNF proteins, and for ORFs detected in other lncRNAs, such as *MALAT1* (Xiao et al. 2024). Notably, consistent with their translatability, the predicted coding potential of these ORFs was significantly higher than the overall scores of all of the eRNA-encoded ORFs, although that of shorter eORFs was lower, reflecting the fact that ORF length is part of the CPAT algorithm (Supp. Table 1). Additionally, all 53 of these transcripts lack introns, as judged by examination of corresponding genomic sequences, consistent with the fact that eRNAs are known to lack introns (Kim et al. 2015). One example, termed ORF5 (see below), is presented as an IGV plot with ribosome footprints and poly (A+) RNA-seq reads, illustrating both the eRNA site identification and ORF detection (Figure 1C). Interestingly, analysis of OpenProt (Leblanc et al. 2024) provided evidence in the form of MS data that at least twelve of the eORF proteins were detectable in cells, even in the absence of Mtr4 KD (Table 3). We note that this list of 53 eORFs is very likely an underestimate of eRNAs with translatable ORFs, as we set a very conservative threshold for filtering, and many eRNAs that are expressed in a tissue/cell type-specific manner may have been missed in our analysis.

### eORF-encoded proteins display many common features

The ten longest ORFs from the above 53 transcripts, ranging in size from 792 to 1332 nt, were selected for further detailed characterization. Expression levels of several of the eORF transcripts were confirmed by RT-qPCR, which matched the previous >2-fold minimum increase in Mtr4-KD cells (Supp. Fig. 2). All ten have coding potentials over 90%, most being close to 100% (Table 1). Notably, among these ten, which we dubbed ORF1-10, ORF4 and ORF8 are found on eRNAs transcribed from the same enhancer but in opposite directions. The others are localized at distinct chromosomal loci throughout the genome (Table 1). The predicted amino acid (aa) sequences of these eORFs revealed some striking features (Table 1). Remarkably, the isoelectric points of the encoded proteins are highly basic, with all ten displaying pIs >11.5. In keeping with this, these proteins contain an unusually high number of arginine residues, 10% to 20% (average of all human proteins is ∼5.53%; calculated from Uniprot database). However, in contrast, the eORF proteins have unusually low numbers of lysine residues, only 0.6% to 3.2% (average is ∼5.81%), indicating that arginine-richness is almost entirely responsible for their basic chemistry. Notably, these properties extend to a large fraction of the 53 high-confidence translated eORFs, with 42 displaying pIs >10 and 40 arginine-richness ≥10% (Supp. Table 1A). Additionally, 15 eRNAs were detected containing ORFs initiating with CTG that match all other criteria used for the 53 ATG-initiated eORFs. Among these, 13 displayed pIs >10, 11 displayed arginine-richness >10%, and 13 had lysine contents <3% (Supp. Table 1B). In light of these observations, we also examined eORF codon frequencies for the six arginine codons, and compared them to the overall frequencies observed across human coding regions. Seven out of the ten largest eORFs had >70% CGN arginine codons (as opposed to AGA/AGG), vs the average of 59%. More notably, CGC and/or CGG codons were especially common compared to the average, with as high as 47% CGC vs 19% average and 44% CGG vs 21% average (Supp. Table 1C). The possible basis for and significance of these properties of eORFs are discussed below.

**Table 1:** ORF and protein characteristics of the ten largest eORF proteins. Properties of the 10 largest ORFs identified by ribosome profiling and used for protein characterization. Chromosome localization of eORFs is indicated. Coding potentials for all sequences were determined by CPAT, and protein chemical characteristics were determined by Expasy ProtParam. Localization predictions were made by DeepLoc 2.1, with results scored above each threshold reported. Proteins sequences tested are highly basic with high numbers of arginine residues, and predicted nuclear and nuclear+cytoplasmic localization patterns.

| **ORF ID** | **Genome Coordinates (hg19)** | **CPAT Coding Potential** | **Protein size (aa)** | **Isoelectric Point (pI)** | **Arg %** | **Lys %** | **Predicted Subcellular Localization (DeepLoc2.1)** |
| --- | --- | --- | --- | --- | --- | --- | --- |
| ORF1 | Chr6: 157695392-157696723 | 1.00 | 443 | 11.69 | 14.7% | 2.3% | Nucleus |
| ORF2 | Chr6: 1604469-1605776 | 1.00 | 435 | 11.82 | 15.2% | 1.4% | Nucleus |
| ORF3 | Chr11: 44748284-44749549 | 1.00 | 421 | 12.05 | 16.6% | 1.2% | Nucleus |
| ORF4 | Chr22: 46466515-46467639 | 1.00 | 374 | 12.16 | 19.0% | 0.8% | Nucleus |
| ORF5 | Chr20: 21486229-21487194 | 0.99 | 321 | 11.74 | 15.9% | 0.6% | Nucleus |
| ORF6 | Chr19: 30363703-30364605 | 0.97 | 300 | 11.75 | 14.0% | 2.0% | Nucleus |
| ORF7 | Chr19: 15334128-15334967 | 0.98 | 279 | 11.88 | 10.0% | 3.2% | Cytoplasm, Nucleus |
| ORF8 | Chr22: 46467763-46468569 | 1.00 | 268 | 11.86 | 16.0% | 2.6% | Nucleus |
| ORF9 | Chr8: 103822496-103823293 | 0.94 | 265 | 12.04 | 15.8% | 0.8% | Nucleus |
| ORF10 | Chr19: 13976122-13976913 | 0.98 | 263 | 12.41 | 16.3% | 1.5% | Nucleus |

We next used computational analyses to predict subcellular localization of the top ten eORFs. For this, we used DeepLoc 2.1, which predicts protein localization using protein language models trained with proteins with known localizations (Odum et al. 2024). This analysis suggested nuclear or cytoplasmic/nuclear localization for all the proteins, consistent with their chemical features (Table 1). Furthermore, DeepLoc 2.1 predicted that all ten eORF proteins are soluble rather than membrane-bound (see Supp Table 2). While these predictions do not provide insight into structure or function, they do provide evidence for similar patterns of subcellular localization, which is examined directly below.

**Table 2:**
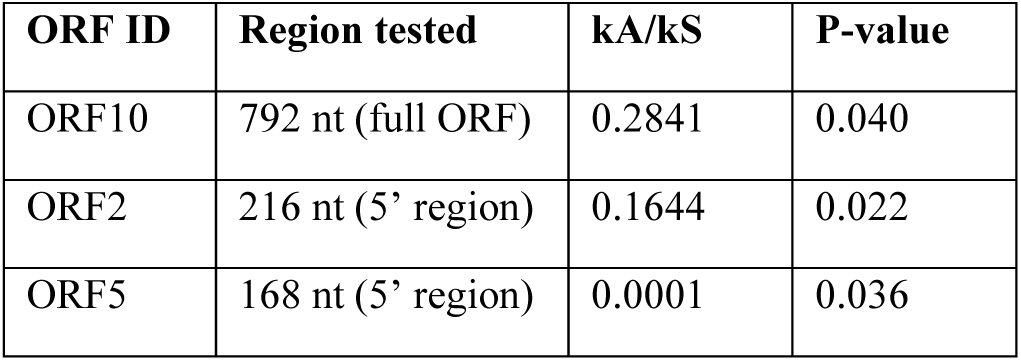
PAML analysis results for significant purifying selection in eORFs. ORFs 2, 5, and 10 have kA/kS values indicating statistically-significant purifying selection across part or all of their sequences. kA/kS values were calculated using PAML with chi-squared p-values determined from likelihood values of these calculations.

We next examined intrinsic features of the top ten eORF proteins, such as ordered vs disordered regions and potential to interact with other proteins, DNA or RNA. For this, we used the DEPICTER2 program, which can predict these and other properties based solely on primary sequence (Basu et al. 2023) (Supp. Fig. 3). The results predicted significant disordered regions present across the eORFs, ranging from 27% to 67% of the amino acid sequence classified as disordered. In many cases, these disordered regions included molecular recognition features (MoRFs), which are regions predicted to undergo disorder-to-order transition through interactions with other proteins (Malhis et al. 2016). These regions constituted up to 34% of the eORF sequences. All the analyzed eORFs were also predicted to have significant protein and RNA binding ability, ranging from 49% to 99% of the residues capable of protein-protein interaction, and 9% to 68% RNA binding. Notably, potential DNA-binding residues were almost entirely absent from these proteins, with only 0% to 3% coverage (Supp. Fig. 3).

### eRNA-encoded proteins are primarily localized to nuclei and appear chromatin-associated

We next examined properties of some of the eORF proteins experimentally, again focusing on the ten largest ORFs. We first designed N-terminal His+FLAG-tagged versions of these proteins and cloned the sequences encoding them into pTWIST-CMV expression vectors (Figure 2A). The resultant plasmids were transfected into HEK293T cells and as a first step expression was examined by Western Blotting (WB) using anti-FLAG antibodies (Figure 2B). All ten proteins were expressed at readily detectable levels and gave rise to bands of the expected size, although there was some variation in accumulation. For example, ORF2 was detected at lower levels than the others, while several, e.g. ORF7, showed decreased accumulation 48 hrs post transfection compared to 24 hrs. Also, several appeared as doublets, notably ORF1, suggestive of posttranslational modification.

**Figure 2:**
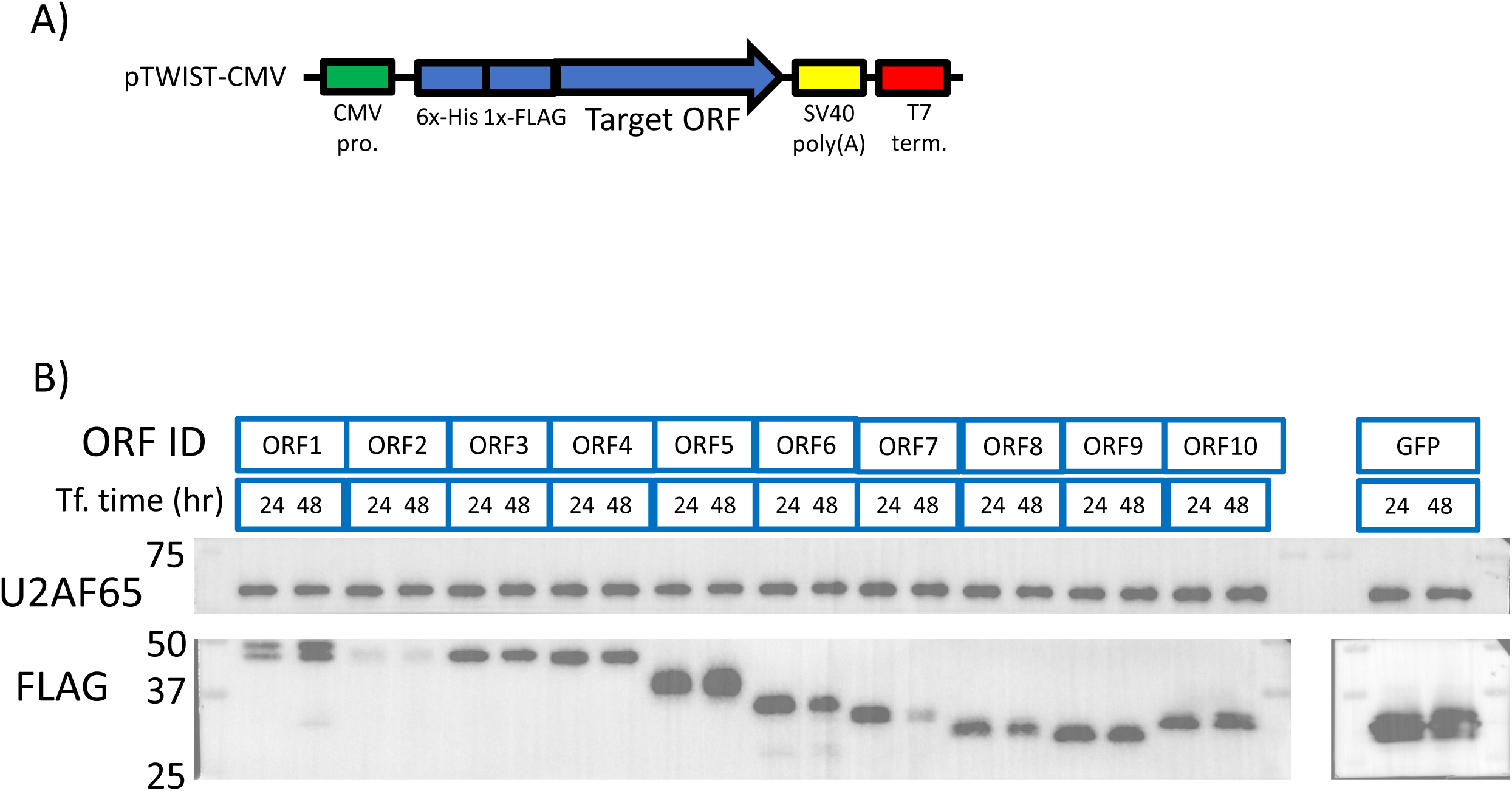
Exogenously expressed eRNA-encoded proteins accumulate in cells. **A** Schematic of the pTWIST-CMV plasmid used for expression of all 10 ORF constructs in mammalian cells. All ORF constructs include a 5’ sequence for a tandem His-FLAG tag for detection and purification methods. Backbone promoter and terminator elements are also shown. **B** Western blotting followed by staining with an anti-FLAG M2 primary antibody and U2AF65 loading control of all 10 eRNA proteins with whole-cell lysates prepared 24 and 48 hours following transfection of HEK293T cells. 3xFLAG GFP was used as a positive control for protein expression under a CMV promoter.

The above expression plasmids were prepared with synthetic cDNAs in which coding sequences were modified to facilitate their synthesis (by lowering local and overall GC content) and, because of the N-terminal tags, translation did not begin at the natural AUG. To verify eORFs could be translated from their natural initiation codon and with authentic coding sequences, we amplified the ORF2 sequence from total cell RNA by RT-PCR using an oligo(dT) primer, and cloned products into pcDNA3 expression vectors. Additionally, to account for possible differences in expression that might arise from different vectors, the N-terminal-tagged ORF2 sequence was also cloned into the same pcDNA3 backbone. Plasmids were transfected into HEK293T cells and expressed proteins analyzed by WB as above. Notably, expression levels of ORF2 derivatives were similar with either tag position, C-terminal or N-terminal (Supp. Fig. 5), providing additional support for the translatability of the eORFs. Only the change of backbone (pTWIST-CMV vs pcDNA3) produced significant expression changes.

We next used immunofluorescent (IF) imaging of transfected HEK293T cells to determine localization of the exogenously expressed proteins. This revealed nuclear localization of all ten ORF proteins, with weaker cytoplasmic signals for some (Figure 3A and Supp. Fig. 3). However, localization of expressed proteins within the nucleus varied. For example, ORF5 and ORF10 were primarily located around the nuclear periphery overlapping with denser chromatin in that region, while ORF2 and ORF8 were mostly found within the nucleoplasm overlapping regions of less-dense chromatin (Figure 3A). Additionally, localization outside of the nucleus varied between proteins: ORF2 appeared in dense clusters while ORF8 was distributed more widely throughout the cytoplasm. ORF10 formed apparent projections from the nucleus with some wider cytoplasmic distribution and nuclear morphology, potentially indicating likely nuclear membrane disruption (Figure 3A). We also performed IF imaging on HEK293T cells expressing either the C- or N-terminal tagged versions of ORF2 (Supp. Fig. 5). Both derivatives displayed the same localization patterns, indicating that position of the tag did not affect localization. While the functional significance of the localization patterns observed remains to be determined, they were largely consistent with the predictions from DeepLoc 2.1 described above, and reflective of the highly charged nature of the proteins.

**Figure 3:**
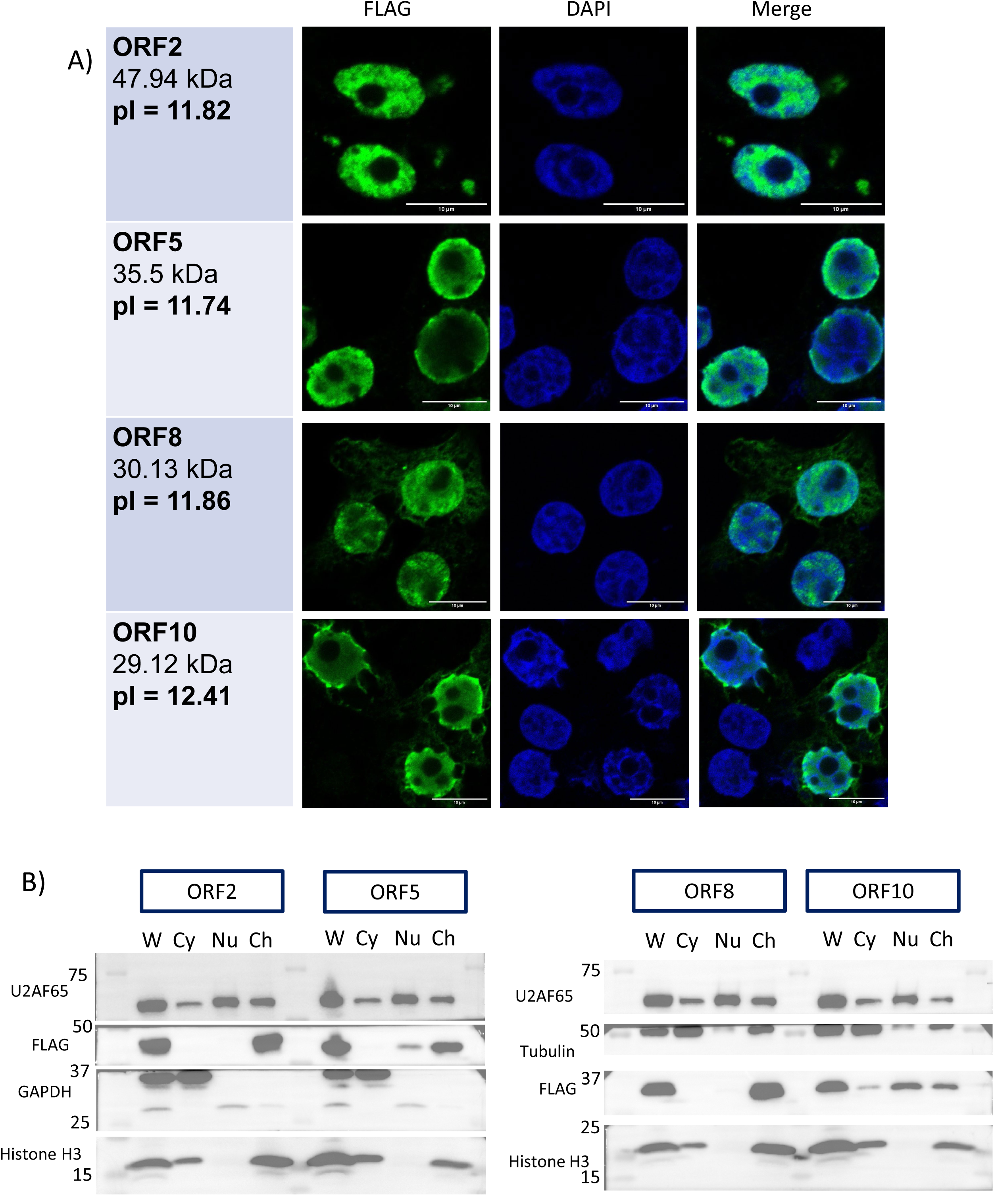
Subcellular localization and fractionation of eRNA-encoded proteins. **A** Immunofluorescent images of HEK293T cells expressing indicated FLAG-tagged eORF proteins. FLAG signal (AF488, green), DAPI (blue) and merged images show localization of four eRNA-encoded proteins, with DAPI showing nuclear DNA. **B** Analysis of eRNA-encoded proteins by subcellular fractionation and western blotting. Fractions tested are whole cell (W), cytoplasm (Cy), nuclear-soluble (Nu), and chromatin (Ch). Fractionation controls used are U2AF65 for the nuclear soluble fraction, Histone H3 for chromatin, and GAPDH or Tubulin for the cytoplasm. Signals match expected localization pattens for those fraction controls.

We next performed biochemical fractionation to examine further the localization of ORFs 1-10. HEK293T cells expressing the tagged proteins were separated into cytoplasm, nucleoplasm (nuclear soluble) and chromatin (nuclear insoluble) fractions as described previously (Ogami et al. 2017; see Methods for details), and proteins detected by WB as above (Figure 3B). Strikingly, the most common pattern for these proteins was localization almost exclusively to the chromatin fraction. The ORF10 protein behaved differently, in that the largest amount was present in the nuclear soluble fraction with smaller amounts in the cytoplasm and chromatin fractions (Figure 3B).

### Immunoprecipitation-mass spectrometry provides evidence for eRNA-encoded protein interactions with RNA processing and DNA damage response complexes

Elucidating the functions of these novel eRNA-encoded proteins will be an important but challenging goal. As a first step, we set out to identify potential interacting proteins for each of the ten eORFs analyzed above. We transfected HEK293T cells with the expression vectors encoding the His-FLAG-tagged derivatives of each protein, prepared cell extracts, and isolated each tagged ORF and associated proteins using two rounds of affinity purification, FLAG-IP followed by TALON beads. All ten purifications were performed with both RNase-treated and RNase-free extracts to identify any RNA-dependent interactions. Co-precipitated proteins were then identified by MS (Methods for details). Tandem MS results for all samples were compared against un-transfected controls to identify significant interactors, defined as exhibiting 2-fold or greater increase in detection over the control in the first (data-independent) stage and ≥10 detection events in the second (data-dependent) stage (Supp. Tables 3). Proteins were also compared between RNase-treated and RNase-free groups to identify RNA-independent, RNA-dependent and, as we somewhat unexpectedly found, RNA-blocked interactions.

Our goal in these experiments was to obtain a general overview of proteins interacting with the ten eORFs we have characterized, but not at this stage to obtain detailed insights into the functional significance of such interactions. For example, given the surprising similarities in the chemical properties and subcellular localization of the proteins, are there also similarities in the interacting proteins? And, especially given the predicted ability of the eORFs to interact with RNA (see above), what fraction of interactions are affected by RNase? As one approach, we used gene ontology (GO) analysis of the MS results to define distinct groups of interacting proteins under both RNase-treated and RNase-free conditions (Supp. Tables 3). In the absence of RNase, all ten eRNA-encoded proteins were found to interact with proteins involved in ribosome structure and function. These included cytosolic small and large ribosomal subunit proteins, which were present in all ten IPs, and others such as pre-rRNA processing proteins and/or mitochondrial ribosomes, which were detected in several (Figure 4A-D; Supp. Table 3B, E, H, J). IPs of ORFs 2, 5, 8 and 10 all included components of the small subunit processome, and ORFs 2, 8 and 10 included other nucleolar proteins. ORF10 also associated with other pre-ribosomal and ribosomal biogenesis proteins, indicating a possible involvement in ribosome processing/assembly. Other RNA-interacting protein complexes were also detected, such as the coding region instability determinant (CRD)-mediated mRNA stability complex. Notably, DNA-interacting protein complexes were also detected, especially those involved in the DNA damage response (DDR) and double-strand break repair (Figure 4A-D; Supp. Table 3B, E, H, J). Other chromatin-interacting proteins, especially histones, were also detected (see also below).

**Figure 4:**
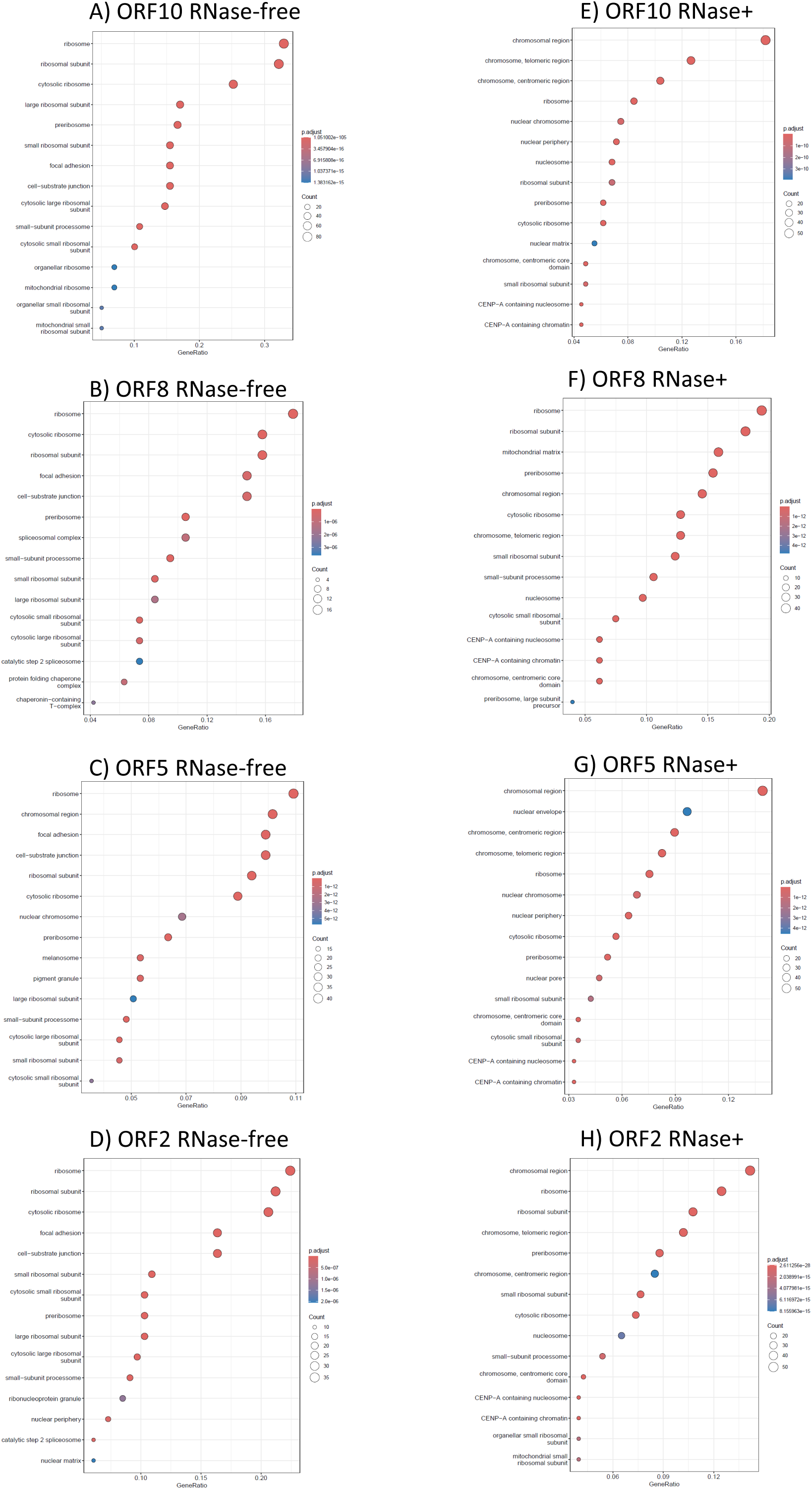
IP-MS proteomics indicate that eRNA-encoded proteins interact with proteins involved in nucleic acid metabolism. Proteins identified by data-independent acquisition on IP-mass spectrometry spectra, grouped by cell component in the Org.Hs.eg.db annotation. The maximum of 15 top significant GO terms are shown with gene ratio relative to the reference list in x-axis, Benjamini-Hochberg adjusted p-values by the indicated color scale, and gene numbers by the point size. Full GO result lists from PANTHER via the Gene Ontoloy Resource, including individual complexes, are provided in Supplementary Table 3. Results for RNase-treated and RNase-free conditions show varying amounts of RNA-mediated vs –independent interactions. The following samples are represented, with both RNase conditions: **A** ORF10, RNase-free. **B** ORF8, RNase-free. **C** ORF5, RNase-free. **D** ORF2, RNase-free. **E** ORF10, RNase-treated. **F** ORF8, RNase-treated. **G** ORF5, RNase-treated. **H** ORF2, RNase-treated.

We next examined how RNase treatment affected the interactions detected in its absence. Ribosomal and rRNA-processing proteins were less represented in the RNase-treated samples, indicating that at least some of these interactions were likely RNA-mediated. These included both cytosolic and mitochondrial ribosomal proteins (Figure 4D, H; Supp. Table 3J, T). Perhaps as expected, interactions with the DDR-related complexes noted above were resistant to RNAse. Notably, some of these complexes appeared to have increased representation in RNase-treated samples, which is discussed further below. Interactions observed with the CRD-mediated mRNA stability complex were also resistant to RNase treatment, suggesting possible direct interactions with component(s) of this complex (Supp. Table 3E, O).

Unexpectedly, several interactions were only detected, or greatly increased, in the RNase-treated IPs. ORF2 showed an especially large change between RNase-treated and untreated conditions, shifting from primarily interacting with rRNA- and mRNA-associated proteins in the RNase-free sample and more with DNA-associated and rRNA processing proteins in the RNase-treated conditions (Figure 4A, E). The DDR complexes noted above were present in RNase-treated ORF 2, 8 and 10 IPs and others (Supp. Table 3L, R, T), despite being absent or less prevalent in their RNase-free counterparts. RNase-treated ORF 8 and 10 IPs included proteins associated with chromosomal telomeric regions, and other RNase-treated IPs also included other telomere-associated proteins (Figure 4G, H; Supp. Table 3R, T). Furthermore, all the histones detected in the RNase-free samples above were present in higher abundance in the RNase-treated samples, and others were only detected in RNase-treated samples. ORF 2, 4, 8, 9 and 10 RNAse-treated IPs also included proteins involved in chromatin binding and/or nucleosome assembly (Figure 4G, H; Supp. Tables 3L, N, R, S, T). The possible significance and basis for proteins appearing in IPs only after RNAse treatment is discussed below.

Analysis of proteins coIPed with all ten ORFs by SDS-PAGE and Coomassie staining revealed patterns largely consistent with those detected by MS (Supp. Fig. 6). Changes in proteins detected indicate the presence of both RNA-mediated and RNA-blocked interactions. One notable difference was the increased presence of proteins ∼15 kDa in all ten RNase-treated samples. This size profile matched that of histones, consistent with their presence in the IP-MS results and confirmed for one, histone H3, by WB (Supp. Fig. 7).

We described above evidence from the OpenProt database indicating that multiple eORF proteins can accumulate to detectable levels in cells. This data represents a genome-wide search for novel ORFs supported by MS from multiple sources, including numerous coIPs from the BioPlex database (Huttlin et al. 2021). Interestingly, one of the OpenProt entries (IP_4463835) we found that matches an eORF (ORF8; Table 3 and Supp. Table 4) was detected in an IP-MS of the putative RNA helicase DHX57, which in turn was among the proteins we detected that coIPed with tagged ORF8. These findings strongly support the physiological significance of the ORF8-DHX57 interaction. Why additional reciprocal interactions were not detected in the OpenProt database is unclear, but could reflect any number of technical and/or biological limitations, e.g., sample preparation, the MS method or data analysis.

**Table 3:** eORF protein matches to OpenProt novel proteins. eRNA ORFs with predicted proteins matching OpenProt novel protein entries are listed. ORF IDs and coordinates match those presented in Table 1 and Supp. Table 1. OpenProt results display the number of studies in which matching peptides were detected by mass spectrometry, the number of unique peptides identified, and the percentage of each eORF represented by detected peptides. Additionally, the two percentages marked with * represent OpenProt entries that cover only a C-terminal region of the matching eORF protein: IP_3415846 covers 34% of the ORF4 sequence, resulting in 4.5% coverage of the full-length eORF, IP_160100 covers 40% of the ORF26 sequence, resulting in 5.3% coverage of the full-length eORF, while IP_4415084 covers 55% of the ORF48 sequence, resulting in 6.0% peptide coverage of full-length eORF. OpenProt in both these cases utilized an internal AUG codon within the eORF sequence, resulting in shorter proteins than we identified. OpenProt entries marked with ** reported peptide matches from MS studies but did not report values for the following columns.

| **ORF ID** | **Coordinates** | **OpenProt ID(s)** | **OpenProt Studies** | **Unique Peptides** | **MS Coverage** |
| --- | --- | --- | --- | --- | --- |
| ORF4 | Chr22: 46466515-46467639 | IP_3415846 | 3 | 2 | 13.2%* |
| ORF8 | Chr22: 46467763-46468569 | IP_4463835 | 5 | 5 | 28.1% |
| ORF12 | Chr13: 107569810-107570502 | XP_047286814.1 | 1 | 1 | 4.8% |
| ORF16 | Chr7: 5467214-5467879 | Q6ZUU7 | 2 | 3 | 18.1% |
| ORF26 | Chr7: 75796016-75796474 | IP_160100 | 1 | 1 | 13.1%* |
| ORF29 | Chr7: 128170418-128170834 | IP_579378** |  |  |  |
| ORF30 | Chr5: 95170588-95171001 | IP_2359134 | 2 | 1 | 6.7% |
| ORF31 | Chr6: 25992638-25993051 | IP_4403603 | 1 | 1 | 8.8% |
| ORF32 | Chr8: 103819070-103819483 | IP_570993 | 2 | 2 | 13.1% |
| ORF35 | ChrX: 135991555-135991959 | Q96NB7 | 1 | 2 | 16.2% |
| ORF48 | Chr19: 33667580-33667915 | IP_4415084 | 5 | 1 | 13.1%* |
| ORF52 | Chr7: 148637904-148638209 | IP_3805337** |  |  |  |

### eRNA ORFs are conserved among primates with homologs from related species providing evidence for recent evolution

We next investigated the evolutionary conservation of the eORFs we have characterized. To address this, we compared sequences of ORFs 1-10 across different species. We extracted likely orthologs for each of these ORFs using NCBI BLASTN against mammalian Refseq genomes, identified any ORF(s) within each ortholog and aligned these sequences using the MAFFT algorithm on Guidance2 (Sela et al. 2015); see Methods for details).

Notably, all ten eORFs have detectable homologs in other primates, but significant homologies, especially complete ORFs, were not detected outside of primates. ORFs 4, 8 and 10, display homology along their entire length across ape and old-world monkey species and encode proteins of similar length and sequence (Figures 5A, B; alignments of all ten ORFs are presented in Figure 5 or Supp. Fig. 8; ORFs 4, 8, 10 in Supp. Fig. 8D, H, J). For example, and most strikingly, ORF10 aa sequence identity ranges from 99.24% with chimpanzee to 91.63% with macaque (Figure 5B). A similar homology pattern was observed for ORF8 across apes and old-world monkeys, and ORFs 3 and 4 among great apes (Supp Fig. 8C, D, H). On the low end of ORFs with complete homologs, ORF9 displays 93.43% identity with bonobo and 85.40% identity with gorilla, while other primate matches (including chimpanzee, orangutan and old-world monkeys) revealed one or more shorter ORFs matching only the N- or C-terminal regions (Supp. Fig. 8I, J). Complete ORFs are not common outside of apes, as most other primate homologs contain one or more smaller ORFs matching the human version. Notably, in all cases any insertions or deletions maintained the reading frame, consistent with selection for the complete protein sequence across these species.

**Figure 5:**
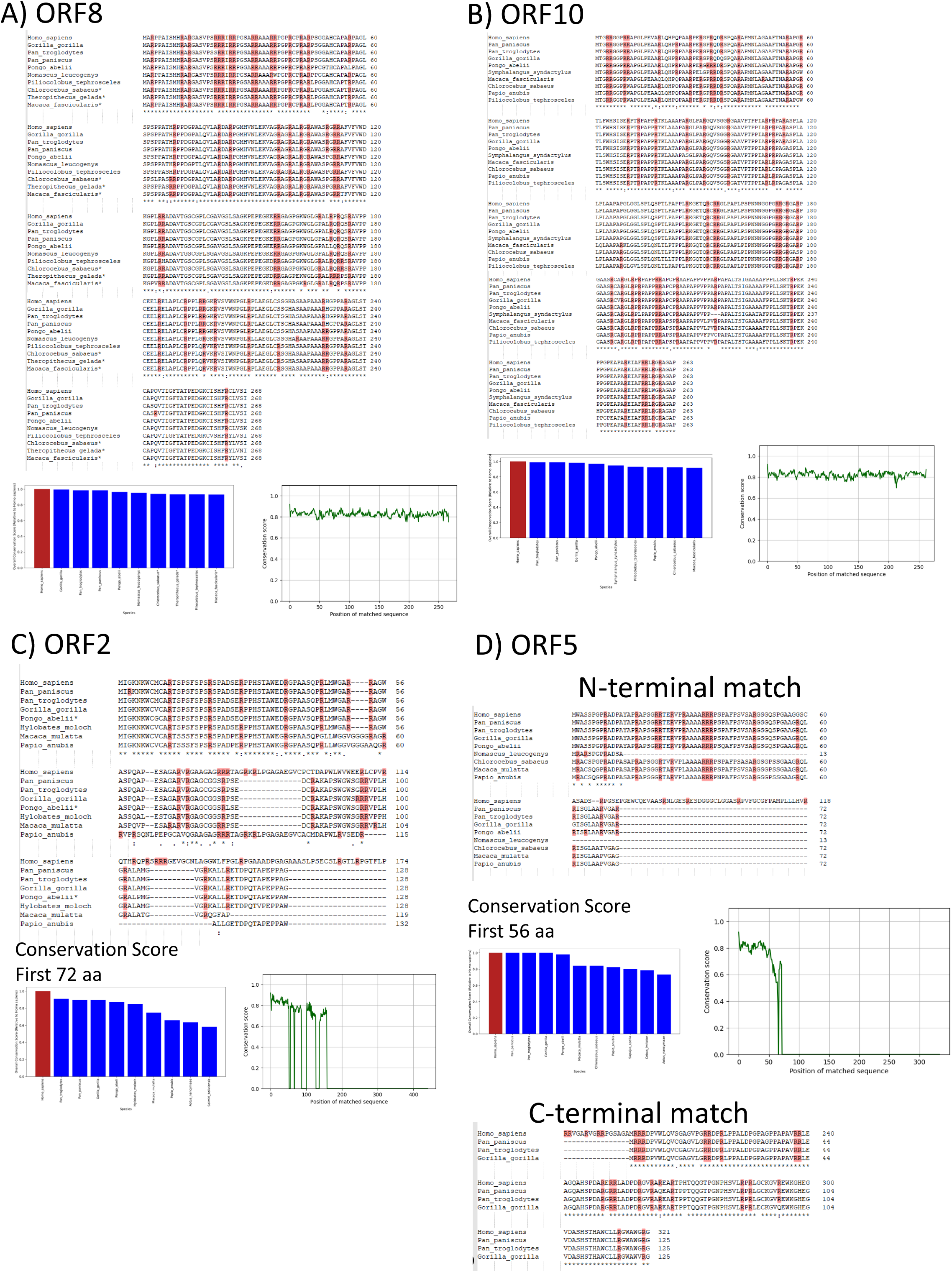
Evolutionary comparison of eORF proteins. Predicted human eORF-encoded proteins were aligned with their homologs in great apes and old-world monkeys (2 or more species among macaques, baboons, and others). Homologous DNA sequences were obtained from NCBI BLASTN, translated using Expasy Translate, and aligned by Clustal Omega. Arginine residues are highlighted. Conservation plots for the amino acid sequences compared to the human versions are presented by percentage of the whole sequence (shown in bar plots) and Jensen-Shannon divergence score per amino acid position (shown in line plots), compared to the whole human sequence unless noted otherwise. All ORF alignments are also presented in Supplementary Figure 7 with additional conservation plots. **A** Human ORF8 and homologs encode 268aa proteins. Alignments and conservation scores are presented for the full sequence. **B** Human ORF10 and homologs encode 263 aa proteins. Alignments and conservation scores are presented for the full sequence. **C** Human ORF2 encodes a 435aa protein while homologs encode shorter proteins around 128aa. Only the N-terminal region of the human protein is shown as no ORF homologs to its C-terminal region were detected. Jensen-Shannon divergence score per aa residue are shown relative to aa position in the entire human sequence; whole-sequence conservation scores are shown for the shared 72aa N-terminal region. **D** Human ORF5 encodes a 321aa protein. Homologs encode shorter proteins:72aa in ape and old-world monkey N-terminal homologs and a 125aa C-terminal chimpanzee, bonobo, and gorilla homolog. Jensen-Shannon divergence score per residue are presented for the N-terminal homolog, whole-sequence conservation scores are presented for the shared 56aa N-terminal region.

Multiple other eORFs show an interesting pattern in which the human protein contains a unique C-terminal extension not found in other species. These human ORFs thus match shorter homologous ORFs in other species with similar or identical sequences at their N-termini, but contain deletions leading to frameshifts that allow extension by utilizing an alternate reading frame not used in the non-human species. For example, ape homologs of human ORF2 contain shorter, 384 nt ORFs while the human version contains a 7 bp deletion, leading to a frameshift that creates a C-terminal extension. The non-human versions thus encode a 128 aa protein that shares a 72 aa N-terminal region with its human counterpart, while the post-frameshift human version is extended to 435 aa in total (Figure 5C). Other examples of C-terminal extensions are present in ORFs 3, 6 and 7, though their homologs in other species are more varied without a clear single mutation leading to the extended human version (Supp. Fig. 8C, F, G).

Another interesting class of human eORFs arose from what can be considered gene fusion. In such cases, two closely linked ORFs are present in different reading frames in non-human primates, with frameshift mutations leading to the longer, fused human-specific ORF. ORF5 is the clearest example of this, with non-human N-terminal homologs encoding a 72 aa peptide, while the human version contains a 1-bp deletion extending the sequence to 321 aa with a shared N-terminal region of 56 aa. But distinguishing eORFs in this class from those described in the previous paragraph, ORF5 homologs from homininae (chimpanzee, bonobo and gorilla) contain a second complete ORF potentially encoding a 125 aa protein that is fully incorporated at the C-terminus of the human sequence by the aforementioned frameshift (Figure 5D). ORFs 1 and 9 display similar patterns of ORF fusion, albeit having a larger set of mutations leading to more varied non-human ORFs, thus lacking a clear single event responsible for the longer human version (Supp. Fig. 8A, I).

For some eORFs, partial conservation with more distantly related species was also observed. ORF5, for example, displays limited similarities with new-world monkeys and lemurs (Supp. Fig 8A). While most eORFs do not display matches outside of primates, ORFs 4, 8 and 10 display short matches across a wider range of eutherian mammals, although these more distant similarities are fragmentary and do not cover the entire eORF sequence (Supp. Fig. 9B). As noted above, the eRNAs encoding ORF4 and ORF8 are transcribed from the same enhancer site but in opposite directions, and interestingly, both have fragmentary matches across a similarly wide range of mammalian species. For example, ORF4 shows unusually strong conservation in the region near its start codon, which is conserved well enough to possibly encode peptides/proteins with a conserved arginine-rich N-terminal region across many mammalian lineages, though this homology does not go beyond ∼60 aa (Supp. Fig. 9B). This may indicate that the enhancer site containing ORFs 4 and 8 is much older than the others and formed distinct, translatable ORFs from the same start site in multiple mammalian lineages. It is notable that such patterns of ORF fusion/extension are features of de novo gene formation, as we discuss below.

The presence of intact ORFs maintained across multiple species indicates a likely selection for the complete protein (as opposed to nucleotide) sequence. To provide more insight into the evolutionary conservation we have observed among these ORFs, we examined the aligned eORF sequences for their ratio of synonymous (kS) vs non-synonymous (kA) mutations using PAML software (Yang 2007; Alvarez-Carretero et al. 2023). Coding sequences exhibit a higher proportion of synonymous mutations (kA/kS<1), indicating stabilizing selection of the aa sequence (Yang 2007; Alvarez-Carretero et al. 2023). Typically, human protein coding genes have kA/kS values of 0.35 or less, and higher scores are uncommon outside of certain specific gene groups (Hurst 2002). We calculated kA/kS ratios within homininae (human, chimpanzee, bonobo, and gorilla) for the eORFs we characterized here (Table 2). Among the ORFs with full-length matching homologs, ORF10 displayed the strongest evidence of purifying selection, with a kA/kS ratio of 0.28 across its entire sequence. For the ORFs with human-specific extensions and fusions, ORF2 and ORF5 showed evidence of strong purifying selection in their shared 5’ regions. Thus, the 5’ 216 bp region of ORF2, encoding the shared 72 aa N-terminal sequence, has a kA/kS ratio of 0.16, but this strong sequence conservation does not extend to the 3’ 168 bp of the non-human ORFs. With ORF5, the 168 bp 5’ region encoding the shared 56 aa N-terminal sequence displays a striking kA/kS ratio of 0.0001, but this strong conservation does not extend to the second 375 bp ORF that is incorporated into the fused human ORF. These low kA/kS values are not maintained when testing a wider group of species (e.g., all primates), indicating that these sequences have only been undergoing purifying selection in recent evolutionary history. Other eORFs analyzed in this study lacked the clear purifying selection of the above three examples. It may be that they are under weaker selection that cannot be detected with significance by PAML, which may in turn be influenced by the higher mutation rate typical of enhancer sites (MacArthur and Brookfield 2004; Villar et al. 2015). However, for the eORFs with low kA/kS values, the sequence conservation we detected points to a significant and evolving role for the encoded proteins in primates, with novel ORFs undergoing strong sequence conservation characteristic of known coding sequences (Alvarez-Carretero et al. 2023).

## Discussion

The experiments reported here provide evidence for translation of eRNA sequences and biological roles for the encoded proteins. We identified a set of human eRNAs containing long open reading frames and provided evidence that they are translated, especially upon stabilization by depletion of the exosome cofactor Mtr4. Strikingly, the eORFs display common features such as highly basic protein chemistry, reflecting high arginine content, and nuclear localization. Identification of interacting proteins revealed proteins involved in DNA repair, ribosome biogenesis and structure, and other functions associated with nucleic acids. Additionally, these eORFs likely formed relatively recently in evolution as homologues were found only in primates, some with extensions that appear unique to humans. All of these observations point to eRNAs as a possible source of novel proteins formed through de novo gene evolution. Below we discuss these and other properties of this novel class of protein-coding transcripts.

eRNAs and mRNAs have been known to share many similarities, but coding potential has not been one of these. For example, similarities between enhancer sites and mRNA promoters are not limited to their interaction with RNAP II and synthesis of capped, polyadenylated transcripts (although eRNAs do not always undergo the later modification) (Sartorelli and Lauberth 2020). Further similarities include the role of transcription factors, both general and sequence-specific, in their activation, and the presence of certain transcription-related histone marks (Core et al. 2014; Henriques et al. 2018; Colbran et al. 2019). Just as enhancers can function similarly to promoters, including some that may have previously been misidentified as promoters (Core et al. 2014), the reverse is also true as core promoters can activate distal genes through promoter-promoter interactions, similar to activation by enhancers (Medina-Rivera et al. 2018). Furthermore, computational predictions of promoters and enhancers have shown that models for enhancer classification are also able to detect promoters but not vice-versa (Colbran et al. 2019). Our findings of eRNA translation indicates a further functional similarity between enhancers and promoters.

A striking feature of the eORFs we characterized is their highly basic chemistry, especially an unusually high percentage of arginine residues. Arginine-rich domains and proteins are not especially common but play roles in several cellular processes, and include ribosomal proteins (Lott et al. 2013), histone proteins and splicing factors (Chandana and Venkatesh 2016). However, these proteins tend to also contain high levels of lysine and/or the arginine-richness is restricted to specific domains, such as RS domains in SR protein splicing factors. Perhaps most similar to the eORF proteins are the two protamines, PRM1 and 2, which are small histone-like proteins that replace histones in sperm, and which are highly enriched in arginine (Balhorn 2007). Intriguingly, this arginine-richness, suggested to provide more stable interaction with DNA (DeRouchey et al. 2013), seems to have evolved in primates and more generally across mammals despite unusually high rates of nonsynonymous mutations (kA/kS = 0.64) compared to most mammalian gene groups (Rooney et al. 2000). This may reflect selection at the level of overall arginine-richness rather than aa sequence itself, perhaps leading to alterations in DNA packing and providing enhanced sperm competitiveness (Rooney and Zhang 1999; Rooney et al. 2000; DeRouchey et al. 2013). Notably, such rapid, potentially ongoing evolution, along with arginine-richness, nuclear localization and possible nucleic acid-related functions, suggests similarities with the eORFs we have described. And finally, PRM1 and PRM2 are tightly linked and located close to multiple active enhancer sites (Barakat et al. 2018), raising the admittedly speculative possibility that protamines may have evolved as eRNA ORFs.

Arginine-rich proteins have been found to be involved in liquid-liquid phase separation (LLPS). This includes for example proteins involved in formation of Cajal bodies, stress granules, and other RNA-containing subcellular bodies (Boeynaems et al. 2017; Sicoli et al. 2024). The chemical differences between arginine and lysine, specifically the greater hydrophobicity and ability to form stronger cation-π interactions with aromatic groups of arginine, leads to lower solubility of arginine-rich proteins, and contributes to their stronger LLPS behavior (Hong et al. 2022). Proteins involved in some of these processes, such as Cajal body formation, were indeed identified by our IP-MS of eORF-associated proteins. Interestingly, another example of a “lncRNA”-encoded protein, RIEP, also contains high arginine and low lysine levels, comparable to those of the eORFs described here (Feng et al. 2023). This raises the possibility that there may be a family of novel arginine-rich, “lncRNA”-encoded proteins, perhaps emerging in evolution only recently, yet to be uncovered.

Arginine-richness of eORFs may correlate with an unrelated property of enhancer regions, which is the presence of CpG islands. Enhancer sites typically display high GC-content and the presence of CpG islands, including orphan CpG islands located far from canonical promoters (Bell and Vertino 2017; Xiong et al. 2018; Pachano et al. 2021). Arginine is encoded by six codons, four of which are CGN. As described above, CGN codons, especially CGC/G, are more common in eORFs than the average for human coding sequences. Interestingly, CGN codon bias has also been observed both in genes with high rates of de novo mutations and in essential genes (Schulze et al. 2020). It thus may be that CpG islands contribute to the arginine-richness that characterizes the majority of the eORFs we have studied. CpG-richness however is not sufficient to give rise to eORFs, as this property of enhancer regions is conserved throughout vertebrates (Pachano et al. 2021), while the eORFs we have described appear to have emerged only in primates. Notably, the sequences encoding the primate-specific RIEP protein mentioned above lie over the rRNA promoter region (Feng et al. 2023), which is also enriched in CpG dinucleotides (Ghoshal et al. 2004). Finally, we note that the interplay between protein coding potential in eRNAs and underlying DNA sequence constraints (CpG islands) is in some ways analogous to that between protein coding potential in general and the presence of regulatory sequences, such as exonic splicing enhancers, in mRNAs (Chen and Manley 2009).

The presence of protein interactions only detectable in RNase-treated samples is notable. Such “RNA-shielded” interactions were recently observed in a large-scale IP-MS of RNA-binding proteins (Street et al. 2023). Some of these may be artifacts from post-lysis interaction facilitated by RNase treatment (Mili and Steitz 2004; Guo et al. 2024). However, others may represent authentic interactions detectable only when RNA-mediated interactions are eliminated. These can be explained by liberation of target proteins from previously inaccessible large RNP complexes or greater binding of target proteins to beads in the absence of interference from such RNP complexes (Street et al. 2023). Further study of individual interactions between target proteins and complexes of interest is necessary to determine the exact nature of these interactions and eliminate possible post-lysis artifacts.

Enhancer activation and eRNA transcription vary widely across cell and tissue types, both between and within species (Sartorelli and Lauberth 2020). Genes encoding proteins that modulate eRNA levels, such as Mtr4, are also differentially expressed between tissue types and in normal vs cancer cells (Fagerberg et al. 2014; Yu et al. 2020; Yu et al. 2023), adding a further layer to possible variation of eRNA levels. Recent studies of eRNAs in embryonic development have also identified changes in their expression patterns across developmental stages (Yu et al. 2024). Additionally, both eRNAs and other lncRNAs interacting with enhancers have been identified as playing a role in the activation of super-enhancers across a wide range of cancer types (Song et al. 2024), potentially extending their role and variation into the genetic dysregulation and reprogramming observed in cancer. Thus an intriguing possibility is that expression of eORFs is differentially regulated, at the levels of both transcription and RNA stability, across a range of cell types, including different tissue types, developmental stages and perhaps cancers.

The eORFs we have described may reflect examples of de novo gene birth. Enhancer sites exhibit a higher rate of mutation compared to typical coding regions, a property that has been linked to the evolution of transcription factor binding sites in their sequences (MacArthur and Brookfield 2004; Villar et al. 2015; Cornejo-Paramo et al. 2024). The presence of ORFs at these sites makes them candidates for rapid formation of novel genes through de novo gene birth and similar processes (Kaessmann 2010; Carvunis et al. 2012; Majic and Payne 2020). One such mechanism, the “transcription first” model, posits that novel genes can form through already transcribed lncRNAs gaining ORFs through mutations, which are translated to produce neutral peptides that can then gain novel functions as a result of mutation and evolutionary selection (Witt et al. 2019; Zheng and Zhao 2022; Peng and Zhao 2024; Zhao et al. 2024). Evidence for such sequence evolution has been observed in human and other primate lncRNAs, with many transcripts producing novel peptides in humans having no translated homologs (Grandchamp et al. 2022; Vakirlis et al. 2022; Broeils et al. 2023).

Orphan genes, which lack clear homology between evolutionary lineages, have been identified as possible results of recent de novo gene evolution. Such genes have been identified as products of both more well-known mechanisms such as duplication-divergence, as well as de novo ORF formation (Cai and Petrov 2010; Tautz and Domazet-Loso 2011; Van Oss and Carvunis 2019). Previous studies of intergenic ORFs in mice and other mammalian species have found mouse-specific ORFs to be more common near enhancer sites, and RNAs containing them were found to be more stable and more likely to be translated than ORFs unrelated to enhancers or promoters (Majic and Payne 2020). The GC-rich nature of enhancers may also be significant, as orphan CpG islands have been found to exhibit evolutionarily recent acquisition of enhancer-specific histone modifications and formation of novel enhancers unique to humans (Kocher et al. 2024). Thus, we suggest that this rapid evolution of enhancer function is not limited to their regulatory roles in transcription, but may also be involved in de novo gene formation. Indeed, the eORFs described here appear to have emerged recently, either being shared among great apes and still mutating relatively rapidly, or with some originating as shorter ORFs undergoing more recent 3’ extensions and fusion events to generate human-specific extended ORFs. This process appears similar to ORF extension patterns predicted by recent computer models of sequence evolution, with short ORFs being either lost or extended into longer ones (Iyengar and Bornberg-Bauer 2023; Lebherz et al. 2024).

In summary, we have uncovered novel, translated ORFs present in enhancer RNAs. The encoded proteins are highly basic and likely function in different aspects of nucleic acid metabolism. Given the rapidly evolving nature of enhancers, the known processes of de novo gene evolution, and the chemical properties of these eRNA-encoded proteins, we propose that enhancers may serve as a source of novel, largely primate or even human-specific regulatory proteins. In any event, our findings both extend the functions of eRNAs and at the same time expand the coding potential of the human genome.

## MATERIALS AND METHODS

### Cell Lines

All RNA-seq and plasmid expression experiments were performed in HEK293T cells grown in DMEM containing 10% fetal bovine serum. Cells were passaged by one minute treatment with 0.25% Trypsin solution at room temperature, with 2-3 day growth periods between passages.

### siRNA transfection

For transfection, cells were seeded in 6-well plates. Cells were transfected with siRNA against Mtr4 or control siRNA against *Renilla* luciferase (siMtr4 and siRLuc, respectively). Cells were grown for 72 hours after transfection. For experiments requiring larger cell numbers, such as ribosome profiling, cells were transferred from 6-well plates to 10-cm dishes 24 hours post-transfection (3 wells per dish for siMtr4-transfected. 2 wells per dish for siRLuc) and grown for the remaining 48 hours.

### Potential ORF prediction in enhancer regions

The predicted enhancer regions were obtained from the ENCODE Project Consortium, followed by extraction of intergenic enhancers. Considering that eRNAs are bidirectionally transcribed from the center of enhancers and typically span ∼ 2 kb (Djebali et al. 2012), we excluded regions where the central positions are located within 2.5 kb of a gene. ORFs of 300 bp or longer and starting with ATG or CTG were detected using orfipy (Singh and Wurtele 2021). Additional coding potential analysis of eRNA ORFs was performed using CPAT (Wang et al. 2013). For each protein sequence, we utilized DEPICTER2 (Basu et al. 2023) to predict disordered residues, MoRF (disorder-to-order transition residues) and binding predictions (protein-, DNA- and RNA-binding). Net charge per residue was estimated using localCIDER (Holehouse et al. 2017). All the results were visualized alongside the distribution of amino acids frequently detected in the intrinsically disordered regions.

### Lysis and Fractionation for Ribosome Profiling

Ribosome profiling protocol was based on methods published by Ingolia and colleagues (Ingolia et al. 2009; Ingolia et al. 2012; Ingolia et al. 2014; Ingolia 2016) and modifications for footprinting lower-expression genes (Couvillion and Churchman 2017). Following 72 hr transfection with siMtr4 or siRLuc, cells were washed 2x with PBS and scraped with 500 uL of polysome lysis buffer. Resuspended cells were disrupted with a dounce homogenizer and centrifuged at 3000 rpm for 5 min at 4**°** C using an Eppendorf 2424 centrifuge to pellet nuclei. Supernatant samples containing cytoplasm were collected and RNA content was measured using a nanodrop spectrophotometer. 450 uL of each sample was treated with Ambion RNaseI (Thermo-Fisher), amount adjusted based on RNA content, for 30 min at room temperature, then treated with 5 uL of Superase*In RNase Inhibitor (Thermo-Fisher). Samples were snap-frozen with liquid nitrogen and stored at -80**°** C.

For sucrose gradient fractionation, thawed RNase-treated cytoplasm samples were centrifuged at 14000 rpm at 4**°** C for 5 min (Eppendorf 2424) to remove any remaining cellular debris. Sucrose gradients were mixed from 10% and 50% sucrose buffers using the gradient mixer of a Biocomp Gradient Station. Samples were loaded on gradients and centrifuged at 39000 xG using a Beckman-Coulter Avanti-E Ultracentrifuge. After centrifugation, gradients were separated using the Biocomp Gradient Fractionator to identify the monosome fractions based on absorbance at 260 nm. Monosome fractions were snap frozen with liquid nitrogen and stored at -80**°** C for RNA extraction.

### Ribosome Footprint RNA Extraction

RNA was extracted from monosome fractions using TRIZOL-chloroform and precipitated with ammonium acetate, isopropanol, and GenElute LPA. Resuspended RNA was treated with DNaseI and extracted a second time by the same method. RNA samples were separated using NOVEX 15% TBE-urea gels. Gels were stained with SYBR-gold and bands corresponding to 27-34 nt were cut out. Gel fragments were crushed and resuspended in water to extract RNA, and RNA was extracted from the liquid using TRIZOL-chloroform. RNA end repair was done using T4 Polynucleotide Kinase (NEB) in the presence of ATP, and end-repaired RNA was extracted using the RNeasy MinElute Cleanup kit with 3.8x volume of ethanol to ensure removal of all ATP from the sample and prevent adapter concatemer formation during library preparation. RNA samples were tested for appropriate footprint length using an Agilent Bioanalyer 2100 and concentration was measured using a Qubit assay.

### RNA-seq library preparation and sequencing

cDNA libraries were prepared using the NEB Next Small RNA Library prep kit. After adapter ligation, rRNA sequences were removed using biotinylated RNA oligos and streptavidin beads (produced by IDT, sequences from Ingolia et al. 2012 (Ingolia et al. 2012)). cDNA libraries were separated from other fragments by NOVEX 6% TBE PAGE, and fragments of around 150 nt (total size of footprint with adapters) were cut out and extracted from the gel. Library size and quality was confirmed using a Bioanalyer 2100 and concentrations were measured using a Qubit assay. Libraries for all samples were pooled for sequencing. Sequencing was performed on an Illumina NextSEQ 500/550 machine at the Columbia Genome Center using a 400m read 1x75 bp sequencing kit. Data was collected and converted to FASTQ format using the Illumina BaseSpace service. Sequencing was performed on 3 replicates per condition (6 total), and repeated a total of 3 times.

### RNA-seq data analysis

RNA-seq reads were uploaded to the Galaxy Europe server (Galaxy 2024) for analysis. Adapter sequences were removed using Cutadapt (Martin 2011) and read length and quality was checked using FastQC (Andrews 2010). Remaining rRNA reads were removed by aligning reads against human rRNA sequences using Bowtie2 (Langmead and Salzberg 2012), with unaligned reads used for further alignment and analysis. Filtered RNA-seq reads were aligned to the human genome hg19 using HISAT2 splice-aware alignment algorithm (Kim et al. 2019). The bigwig files were generated using deeptools (Ramirez et al. 2016), and the read coverage was visualized in IGV (Robinson et al. 2011). PAS-seq data was obtained from GSE267857 (Soles et al. 2024). Transcript sequences were assembled from reads using Stringtie guided with a transcript database containing the human transcriptome and all predicted eRNA sequences. Read counts from Stringtie (Pertea et al. 2015) were quantified and compared using DESeq2 (Love et al. 2014).

ORFs were identified and their translation was verified using RiboTricer (Choudhary et al. 2020). Reads were separated by length and lengths with the highest read periodicity values across published CDS sequences were selected for analysis of putative eRNA ORFs. Novel ORFs in sequences labeled as noncoding were identified automatically by RiboTricer, and those with a minimum of three codons occupied by reads and periodicity matching the minimum for human ORFs, phase score 0.44, were labeled as coding. Putative eRNA ORFs with a significant increase of 2-fold or greater in read numbers after Mtr4 knockdown, as calculated by DESeq2, and passing the read periodicity test in RiboTricer were identified as likely coding, and the largest ORFs in this group were selected for protein characterization testing.

eORFs were compared to OpenProt novel ORFs, using bedtools intersect (Quinlan and Hall 2010) and BLAST+ (Camacho et al. 2009) on the Galaxy Europe server. OpenProt annotations were downloaded in BED and protein FASTA formats from https://www.openprot.org/downloads. Protein sequences were also searched against a custom BLAST database built from the OpenProt protein FASTA using BLAST+ protein mode. For eORFs with multiple matches, the largest complete match was reported.

### eRNA ORF sequence evolution analysis

ORF sequences selected for protein characterization were analyzed for their sequence similarity with homologous sequences in related species. Homologous sequences were identified using NCBI BLASTN search against all recent genomes in the database. For sequences where homologous ORFs exhibited a different length than the human version (typically shorter in other species), only the coding sequence was selected for further analysis. Homologous sequences were aligned using the MAFFT algorithm in the Guidance2 server (Sela et al. 2015). Phylogenetic trees for represented species were downloaded in newick format from the Timetree server (Kumar et al. 2022). kA/kS analysis of sequences was performed using PAML through the codeML program on the Galaxy Europe server. Visual representations of aligned sequences were obtained using NCBI MSA Viewer and Clustal Omega. The per-residue conservation scores (Jensen-Shannon divergence score) were estimated using score_conservation.py (Capra and Singh 2007) using human sequence as a reference, BLOSUM62 as a background and a Clustal Omega alignment files as an input.

### eRNA ORF Plasmid Design, Growth and Expression

Plasmid insert sequences were designed from the largest coding ORFs identified by ribosome profiling. ORFs were tagged with a 5’ his-tag and FLAG-tag for immunofluorescence and immunoprecipitation experiments. Insert sequences were synthesized and cloned into plasmids through TWIST Bioscience, using the pTWIST-CMV plasmid backbone for expression in mammalian cells. Some codon optimization was required for compatibility with TWIST gene synthesis methods, but the aa sequences were unchanged. Plasmids propagated in *E. coli* and extracted using the IBI I-Blue Plasmid Midi-Prep kit. Plasmid sequences and quality were confirmed by sequencing through Gene-wiz.

cDNA amplification was performed using PCR with Phusion DNA polymerase and primers matching the 5’ and 3’ ends of target eORFs, including a C-terminal FLAG and his-tag sequence in the 3’ primers. Amplified and tagged eORFs were cloned into a pcDNA3 plasmid vector under a CMV promoter. Additionally, to control for differences in protein expression levels between plasmids, N-terminal tagged constructs were also cloned out of the pTWIST-CMV vector into pcDNA3. Vector control plasmids were the empty pcDNA3 and a pTWIST-CMV containing only the his- and FLAG-tag sequences under the CMV promoter.

Small-scale plasmid transfection in 6-well or 12-well plates was done using Lipofectamine 2000. Transfections were done with 1 ug of plasmid DNA and 4 uL of Lipofectamine 2000 solution per well in 6-well plates, or half of each amount respectively for 12-well plates. Transfections were done for 24 or 48 hours. For larger-scale expression, PEI-MAX was used instead. Cells in 10-cm dishes were transfected with 15 ug of plasmid and 180 uL of PEI-MAX solution.

### Immunofluorescent Localization of Expressed eRNA Proteins

HEK293T cells were seeded on gelatin-coated slides in 6-well or 12-well plates for plasmid transfection using Lipofectamine 2000 (ThermoFisher). Transfected cell slides were collected after 24 or 48 hours. Cells were washed with PBS on ice, fixed with 4% paraformaldehyde, permeabilized with 0.1% Triton X-100, and blocked with Pierce Protein-free Blocking Buffer. Cells were stained with Anti-FLAG M2 antibody (Millipore) diluted 1:500 in Pierce Blocking buffer for 1 hour at room temperature, then stained with Anti-mouse Alexafluor 488 secondary antibody diluted 1:500 in Pierce Blocking buffer for 1 hour at room temperature. Stained cell slides were mounted using Abcam DAPI Fluoroshield Mounting Medium and sealed with transparent nail polish.

Fluorescent imaging was performed using a Zeiss LSM800 confocal microscope with a 63x oil immersion lens, and images were collected using the ZEN Blue software. Images were processed using ImageJ FIJI software.

### Immunoprecipitation of Expressed Proteins for Mass Spectrometry and SDS-PAGE

HEK293T cells were seeded on 10-cm dishes and transfected with plasmids using PEI-MAX. Four plates were prepared per ORF sample. 24 hours post-transfection, cells were scraped with lysis buffer and disrupted my sonication (10 pulses, repeated 3 times per sample). For RNase-treated samples, cell lysates were treated with RNaseA for 45 min at 4**°** C on a rotator. For RNase-free samples, lysates were rotated at 4**°** C for 45 min without RNaseA. Samples were then centrifuged for 10 min at 14,000 RPM in an Eppendorf 2424 centrifuge at 4**°** C to clarify the lysates. Anti-DYDDDDK primary antibody (Cell Signaling Technologies) was added to all samples, which were left to rotate at 4**°** C overnight. The following day, samples were mixed with Pierce Protein A/G magnetic beads, incubated for 3 hours at 4**°** C, and washed to remove unbound protein. FLAG-tagged protein complexes were eluted using 3xFLAG Peptide buffer and mixed with equilibrated TALON Metal Affinity Resin (Takara) beads and incubated at 4**°** C overnight on a rotator. The following day, samples were washed and aliquoted for either mass spectrometry sample preparation or SDS-PAGE.

Samples for SDS-PAGE were treated with 2x sample buffer containing 10% beta-mercaptoethanol directly on the resin beads. Protein expression and immunoprecipitation was confirmed using western blotting, gels used were NOVEX 10-20% Tris-Glycine Plus gradient gels, proteins transferred to PVDF membranes using a TransBlot Turbo semi-dry transfer system (Bio-Rad). Membranes were stained for FLAG-tagged proteins using mouse anti-FLAG M2 primary antibody and anti-mouse HRP secondary antibody. Other proteins isolated by IP were detected using Coomassie staining. The same gel format was used as for Western blotting, and gels were stained with BioRad QC Colloidal Coomassie Stain overnight and destained with water.

### Data-Independent Analysis Mass Spectrometry

The TALON resin was incubated in 80µL of buffer (2M urea, 50mM Tris, pH 7.5, 1 mM dithiothreitol, and 5 µg/mL trypsin (Promega: V511C)) for 1 hour at 25° C and 1000 rpm (Eppendorf Thermomixer C shaker) to partially digest the proteins off the resin. The supernatant containing the partially digested proteins was collected, constituting an initial eluate. Two additional 60 uL washes (2M urea, 50mM Tris, pH 7.5) were performed on the resin; the supernatant of each wash was collected and added to the initial eluate to maximize yield. The full eluate was clarified by spinning at 5000g (Sorvall Legend Micro 21).

Half of the partially digested proteins were further reduced with 5 mM DTT for 30 minutes at 25° C and 1000 rpm, then alkylated in the dark with 10 mM iodoacetamide for 45 minutes at 25° C and 1000 rpm. Samples were then digested with 0.5 µg of trypsin overnight (18 hours at 25° C and 700 rpm) and subsequently acidified with formic acid (for a final formic acid concentration of 1% and a sample pH of ≤ 3). The acidified tryptic peptides then underwent a C18 StageTip clean-up as described (Rappsilber et al. 2007). Peptides on the C18 column were eluted with 60 µL of 60% acetonitrile/0.2% formic acid and dried in a vacuum concentrator. Following evaporation, the peptides were reconstituted in 13 µL of 3% acetonitrile/0.2% formic acid for injection onto the mass spectrometer for LC-MS/MS analysis.

### LC-MS/MS analysis on a Q-Exactive HF

About 1 μg of total peptides were analyzed on a Waters M-Class UPLC using a 15 cm IonOpticks C18 1.7 µm column coupled to a benchtop Thermo Fisher Scientific Orbitrap Q Exactive HF mass spectrometer. Peptides were separated at a flow rate of 400 nL/min with a 90-minute gradient, including sample loading and column equilibration times. Data were acquired in data-dependent mode using Xcalibur software; the 12 most intense peaks in each cycle were selected for MS2 analysis. MS1 spectra were measured with a resolution of 120,000, an AGC target of 3e6 and a scan range from 300 to 1800 m/z. MS2 spectra were measured with a resolution of 15,000, an AGC target of 1e5, a scan range from 200–2000 m/z, and an isolation window width of 1.6 m/z.

Raw data were searched against the human proteome (UP000005640) with MaxQuant (v2.6.3.0). To determine protein enrichment within a sample, the ppm value of a protein’s MS1 intensity value was calculated. This was done by dividing a protein’s intensity by the sum of all protein intensities in the respective sample, then multiplying the resulting fractional value by 1,000,000. After that, a pseudocount of +1 was applied before the ppm values were log2-transformed. The log2 values of a sample, relative to the log2 values of a negative control, were subsequently used as an indicator for protein enrichment. The “MS/MS count” – a semiquantitative measure of the amount of peptides identified in MS/MS that match a given protein – was also used as a filter for enrichment with a cutoff of “MS/MS count” ≥10. The significant interactor proteins identified by IP-MS in each condition were subjected to gene ontology analysis using clusterProfiler (Xu et al. 2024) using org.Hs.eg.db (Carlson 2019) as a reference; and using the Gene Ontology Resource (Ashburner et al. 2000; Gene Ontology et al. 2023) GO Enrichment Analysis tool (Thomas et al. 2022) using all proteins detected by IP-MS as a reference.

### Subcellular Fractionation

Fractionation was performed using the method described (Ogami et al. 2017). HEK293T cells were seeded in 10-cm dishes and transfected with plasmids using PEI-MAX, with one plate prepared per sample. 24 hours post-transfection, cells were washed with PBS on ice and scraped with PBS. Cells were pelleted by centrifugation at 100 xG at 4° C, resuspended in hypotonic swelling buffer, and incubated on ice for 15 min. After swelling, half of the cell suspension was reserved as a whole-cell sample and the remaining half was treated with NP-40 at 0.1% final concentration. Cell suspensions and detergent were gently mixed by tapping the tubes and centrifuged at 6000 rpm at 4° C for 5 min (Eppendorf 2424). Supernatants were saved as cytoplasm fractions and pellets were resuspended in detergent-free swelling buffer and centrifuged again as a wash step. Pellets were resuspended in 50% glycerol buffer and then mixed 1:1 with 2M urea nuclear lysis buffer. Samples were pulse-vortexed 3 times, incubated for 1 min on ice, and centrifuged at 14000 rpm at 4° C for 2 min (Eppendorf 2424). Supernatants were saved as nuclear soluble fractions and pellets were washed once with 1:1 glycerol and nuclear lysis buffer mixture. Pellets were then dissolved in water as chromatin fractions. Samples were run together for SDS-PAGE and western blotting. Fractionation was tested using GAPDH or Tubulin as cytoplasm markers, U2AF65 as a nuclear soluble fraction marker, and histone H3 as a chromatin marker. FLAG-tagged proteins were detected using mouse anti-FLAG M2 primary antibody and anti-mouse HRP secondary antibody.

## ACKNOWLEDGEMENTS

This work was supported by NIH grant R35 GM118136 to J.L.M. M.J. was funded by the NIH (R35GM128802; R01AG071869 and R01HG012216) and NSF (Award 2224211). K. O. was funded by JSPS Grant-in-Aid for Scientific Research (C) (JP22K06925) and TAKEDA Science foundation. We would like to thank current and former members of the Manley lab who helped with this project at various points: Pedro Bak-Gordon, Jian Zhang, Claire Sattler, Thomas Yao, Minu Seong, Takahiro Seimiya, Lizhi Liu, Patricia Richard, Albert Tsai and Shuang Feng. We also thank Blaine Loughlin (Columbia University) for help with the tagged eORF protein construct plasmids. We would like to thank Stirling Churchman and Iliana Soto (Harvard Medical School) for help developing the ribosome profiling protocol used here. We would like to thank other current and former members of the Jovanovic Lab, especially Lena Street, Jenny Kim and Abdurrahman Keskin, for their help with ribosome profiling and mass spectrometry experiments. We also thank Naima Okami and Flora Borne (Columbia University) for their help with evolutionary analysis using PAML, and Brian Heubel and Asja Guzman (Columbia University) for help with confocal microscopy. We also thank Laura Landweber (Columbia University) for multiple thoughtful suggestions.

Data analysis was carried out on the Galaxy Europe server. The authors acknowledge the support of the Freiburg Galaxy Team: Person X and Björn Grüning, Bioinformatics, University of Freiburg (Germany) funded by the German Federal Ministry of Education and Research BMBF grant 031 A538A de.NBI-RBC and the Ministry of Science, Research and the Arts Baden-Württemberg (MWK) within the framework of LIBIS/de.NBI Freiburg. Sucrose gradient fractionation was performed at the Columbia Precision Biomolecular Characterization Facility. RNA sequencing was performed at the JP Sulzberger Columbia Genome Center. RNA-seq library Bioanalyzer and Qubit analyses were performed by the Columbia Herbert Irving Comprehensive Cancer Center Molecular Pathology Shared Resource, Molecular Biology Services.

## Author Contributions

P.A.V., K.O., M.J., and J.L.M. designed the research. P.A.V., K.O., E.V., R.K.K., and M.J. performed the research. P.A.V., K.O., E.V., and R.K.K. analyzed data. P.A.V, K.O., E.V., R.K.K., M.J., and J.L.M. wrote the manuscript.

## SUPPLEMENTARY TABLES

**Supplementary Tables 1:** eRNA-encoded ORFs with canonical (ATG) and non-canonical (CTG) identified by ribosome profiling starts share common patterns of basic chemistry and arginine content. **A** 53 ORFs starting with ATG and ≥300 nt identified by ribosome profiling. All ORFs include coordinates, protein sizes, protein isoelectric points, arginine content, and lysine content. **B** 15 ORFs starting with CTG identified by the same method as in Supp Table 1A. **C** Arginine codon percentages for the 10 largest eORFs, encoding the same ORF1-10 proteins discussed in other figures.

**Supplementary Tables 2:** eORF protein DeepLoc 2.1 analysis shows localization predictions. **A** DeepLoc 2.1 predictions, including subcellular localization scores, are presented for all 10 eORF proteins tested along with the score threshold for each localization category. Threshold-passing localization predictions are underlined and also presented in table in Figure 1D. **B** DeepLoc 2.1 solubility prediction scores are presented for all 10 eORF proteins tested.

| ORF ID | Cytoplasm | Nucleus | Extracellular | Cell membrane | Mitochondrion | Endoplasmic reticulum | Lysosome/Vacuole | Golgi apparatus | Peroxisome | Predicted localization |
| --- | --- | --- | --- | --- | --- | --- | --- | --- | --- | --- |
| ORF1 | 0.4024 | <u>0.6458</u> | 0.3093 | 0.1251 | 0.3804 | 0.1135 | 0.0367 | 0.1225 | 0.0223 | Nucleus |
| ORF2 | 0.3989 | <u>0.6463</u> | 0.401 | 0.11 | 0.3416 | 0.1686 | 0.0395 | 0.1212 | 0.0152 | Nucleus |
| ORF3 | 0.3886 | <u>0.6462</u> | 0.2304 | 0.1337 | 0.2903 | 0.1149 | 0.0774 | 0.1105 | 0.0052 | Nucleus |
| ORF4 | 0.361 | <u>0.602</u> | 0.5094 | 0.1015 | 0.2561 | 0.0983 | 0.0642 | 0.1718 | 0.0134 | Nucleus |
| ORF5 | 0.4707 | <u>0.5338</u> | 0.3238 | 0.1355 | 0.4127 | 0.1684 | 0.0769 | 0.1589 | 0.0044 | Nucleus |
| ORF6 | 0.4171 | <u>0.5021</u> | 0.297 | 0.1718 | 0.4732 | 0.1999 | 0.0589 | 0.1789 | 0.0237 | Nucleus |
| ORF7 | <u>0.4913</u> | <u>0.5031</u> | 0.283 | 0.2137 | 0.3893 | 0.1983 | 0.0738 | 0.1551 | 0.0706 | Cytoplasm, Nucleus |
| ORF8 | 0.4562 | <u>0.6253</u> | 0.3675 | 0.1145 | 0.3763 | 0.1082 | 0.0689 | 0.2073 | 0.011 | Nucleus |
| ORF9 | 0.3939 | <u>0.5479</u> | 0.3586 | 0.1249 | 0.3452 | 0.1989 | 0.0924 | 0.193 | 0.0158 | Nucleus |
| ORF10 | 0.4426 | <u>0.5901</u> | 0.2568 | 0.0602 | 0.1862 | 0.1092 | 0.0672 | 0.135 | 0.021 | Nucleus |
| Thresholds | 0.4761 | 0.5014 | 0.6173 | 0.5646 | 0.622 | 0.609 | 0.5848 | 0.6494 | 0.7364 |  |

B)
| ORF ID | Peripheral | Transmembrane | Lipid anchor | Soluble | Predicted membrane association |
| --- | --- | --- | --- | --- | --- |
| ORF1 | 0.223 | 0.334 | 0.21 | <u>0.82</u> | Soluble |
| ORF2 | 0.322 | 0.207 | 0.28 | <u>0.832</u> | Soluble |
| ORF3 | 0.353 | 0.269 | 0.34 | <u>0.793</u> | Soluble |
| ORF4 | 0.358 | 0.197 | 0.253 | <u>0.844</u> | Soluble |
| ORF5 | 0.333 | 0.256 | 0.381 | <u>0.807</u> | Soluble |
| ORF6 | 0.235 | 0.27 | 0.45 | <u>0.805</u> | Soluble |
| ORF7 | 0.347 | 0.289 | 0.266 | <u>0.751</u> | Soluble |
| ORF8 | 0.197 | 0.372 | 0.297 | <u>0.784</u> | Soluble |
| ORF9 | 0.303 | 0.259 | 0.265 | <u>0.804</u> | Soluble |
| ORF10 | 0.398 | 0.128 | 0.218 | <u>0.833</u> | Soluble |
| Thresholds | 0.6 | 0.51 | 0.82 | 0.5 |  |

**Supplementary Table 3:** Gene Ontology results for all 10 eRNA proteins under RNase-free and RNase-treated conditions. All Gene Ontology results were obtained from significant interactor proteins identified by IP-MS in each condition, and results were analyzed with PANTHER GO database. All results include Biological Process, Cellular Component, and Molecular Function group categories. **A**-**J** present ORF1-10 RNase-free samples, labeled as ORF#_N respectively. **K**-**T** present ORF1-10 RNase-treated samples, labeled as ORF#_P respectively.

**Supplementary Table 4:** eRNA ORF matching OpenProt-identified novel proteins detected in published MS data. eORF proteins and their corresponding OpenProt entries from Table 3 are listed with bait proteins of BioPlex IP-MS datasets containing the detected peptides. DHX57 (marked with *) was additionally detected as an interactor with ORF8 and other eORF proteins in IP-MS experiments.

| **eORF ID** | **OpenProt ID** | **BioPlex Interactor** |
| --- | --- | --- |
| ORF4 | IP_3415846 | CNKSR3 |
| ORF8 | IP_4463835 | CDK3 |
|  |  | LPIN3 |
|  |  | DHX57* |
| ORF16 | Q6ZUU7 | SLC25A11 |
|  |  | BZW2 |
|  |  | RRS1 |
|  |  | IL17RE |
| ORF29 | IP_579378 | RNF31 |
| ORF30 | IP_2359134 | C22orf23 |
| ORF32 | IP_570993 | MYL6 |
|  |  | LMBRD2 |
| ORF35 | Q96NB7 | UXT |
|  |  | ZNF232 |
|  |  | CSNK1E |
| ORF48 | IP_4415084 | CAP1 |
| ORF52 | IP_3805337 | TMEM206 |

## SUPPLEMENTARY FIGURES

**Supplementary Figure 1:**
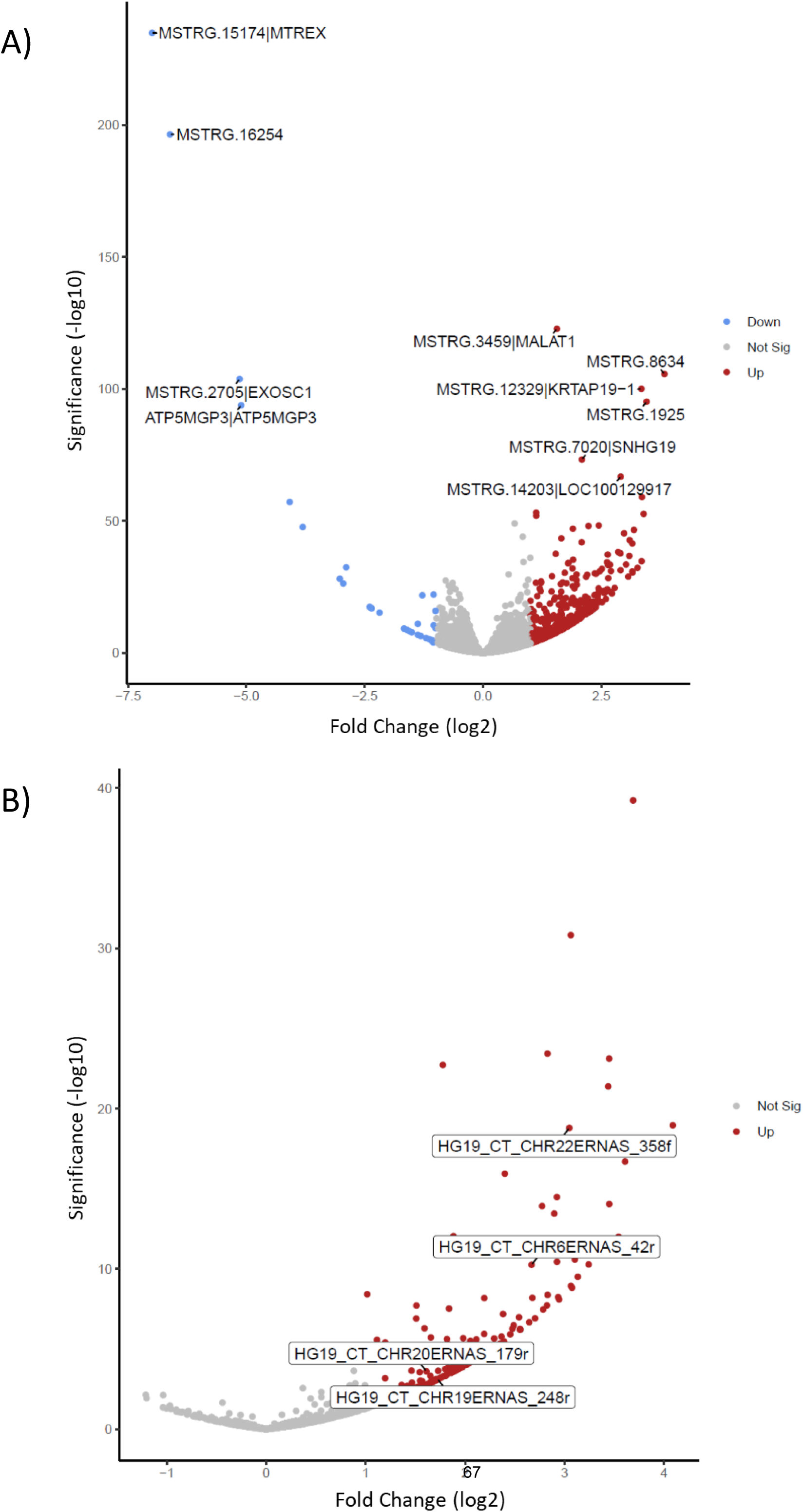
Volcano plots of ribosome profiling data analyzed by StringTie and DESeq2 pipeline. Differential expression results from ribosome profiling experiments comparing Mtr4-KD and siRNA control samples are represented by volcano plots. Fold-change of 2x or greater and adjusted p-value of 0.05, as calculated by DESeq2, are used as cutoffs for significant increase or decrease in expression. **A)** Results for all transcripts assembled by StringTie (starting with MSTRG). These include sequences from published coding and non-coding RNAs, along with novel transcripts detected by the program. **B)** Results for all sequences matching predicted eRNAs. Four marked examples are eRNAs containing ORFs presented in main figures: CHR6ERNAS_42r = ORF2, CHR20ERNAS_179r = ORF5, CHR22ERNAS_358f = ORF8, CHR19ERNAS_248r = ORF10.

**Supplementary Figure 2:**
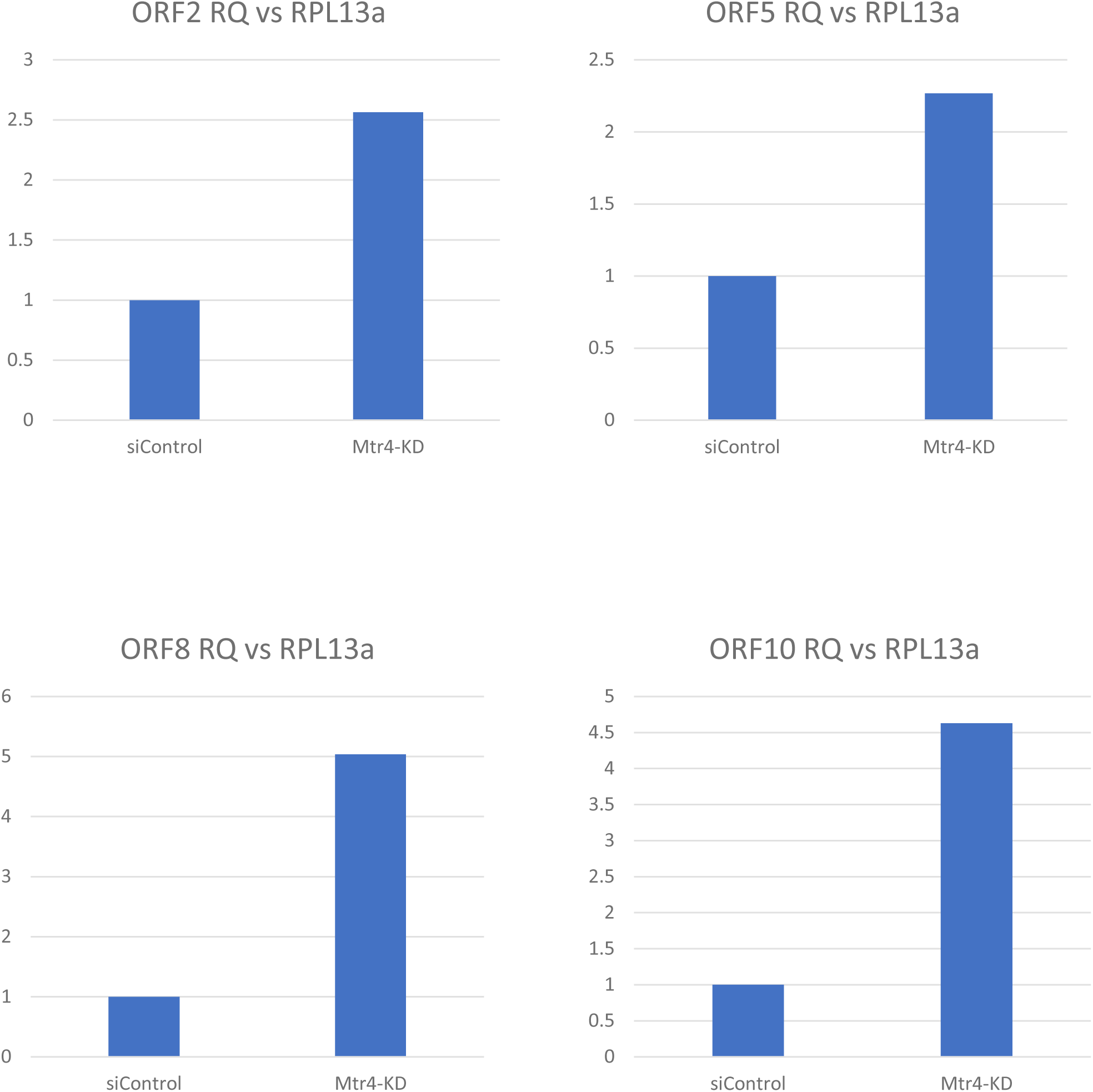
RT-qPCR analysis of eORF transcripts. All samples presented were oligo-dT cDNA of whole cell RNA extracts from Mtr4-KD and siRNA control cells. All results are presented as relative quantification values vs RPL13a control. Targets tested are ORF2, 5, 8, and 10.

**Supplementary Figure 3:**
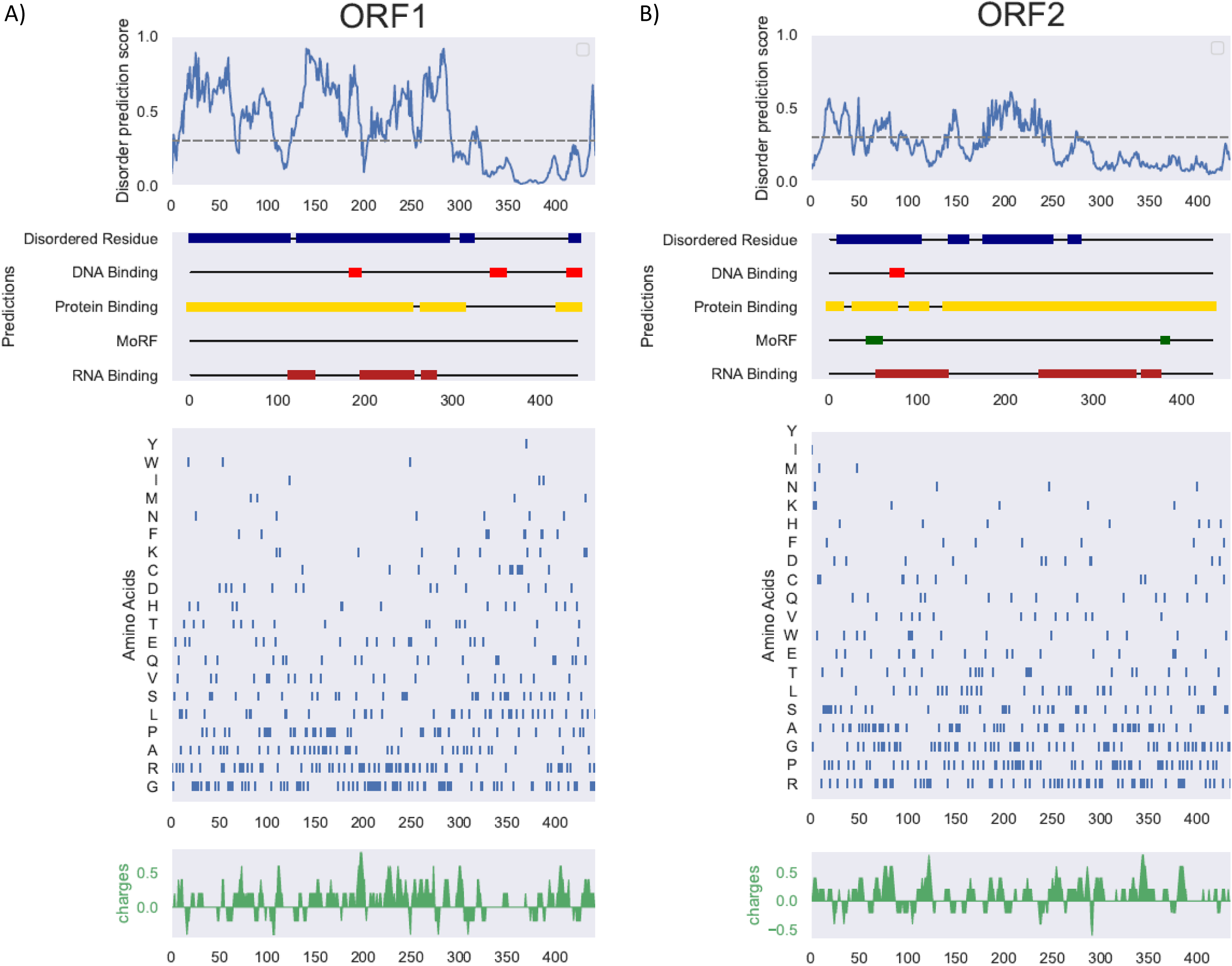
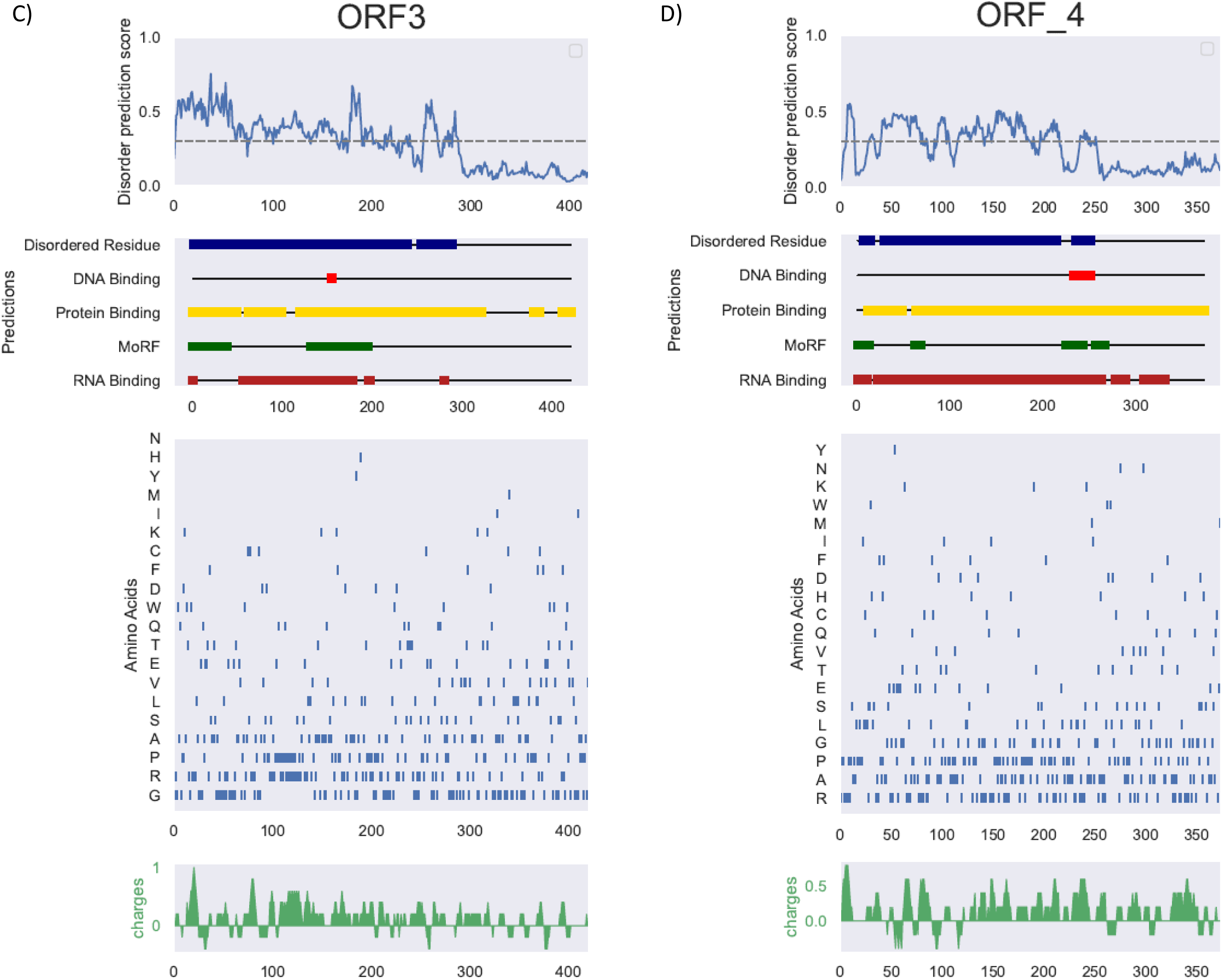
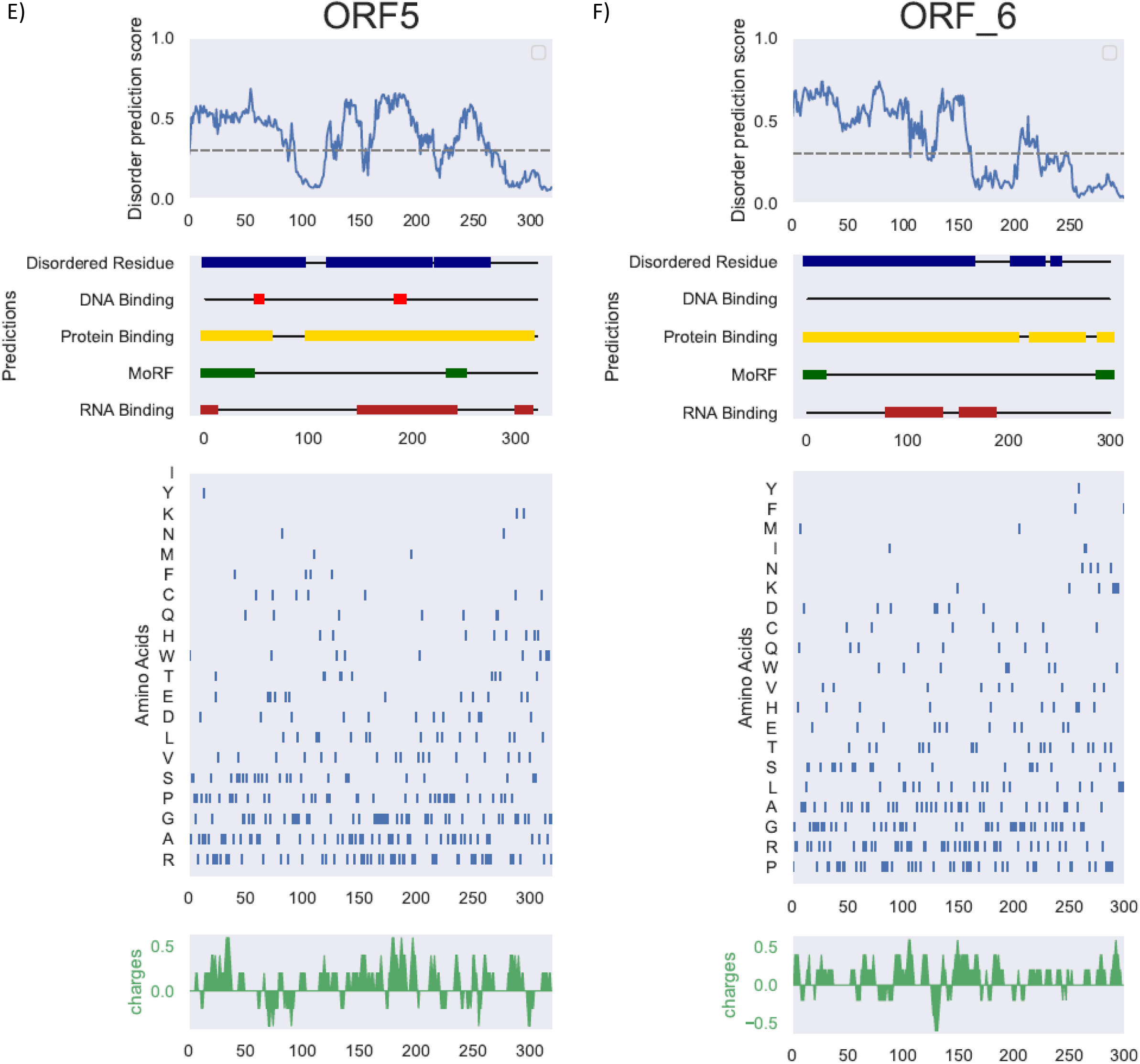
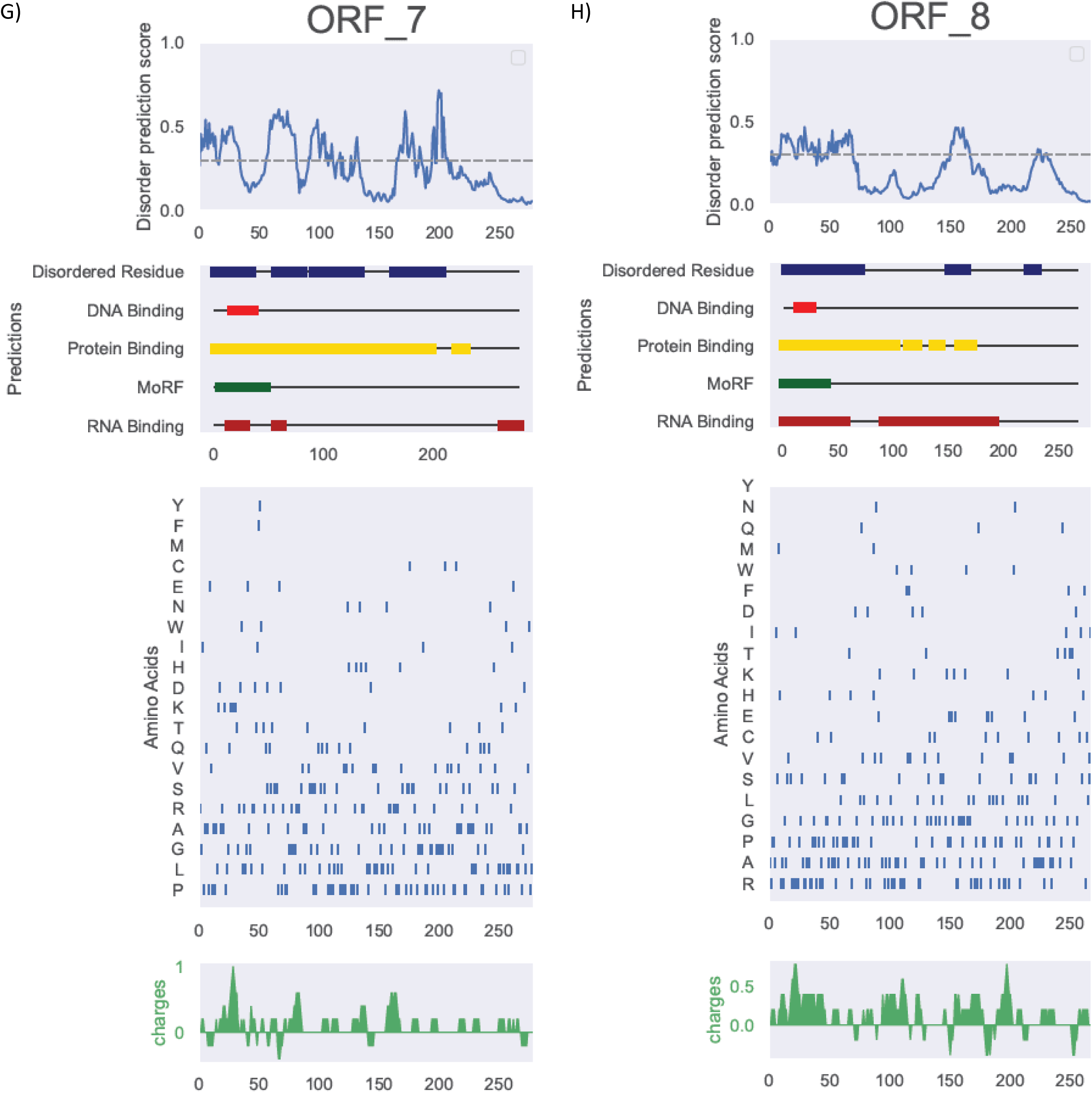
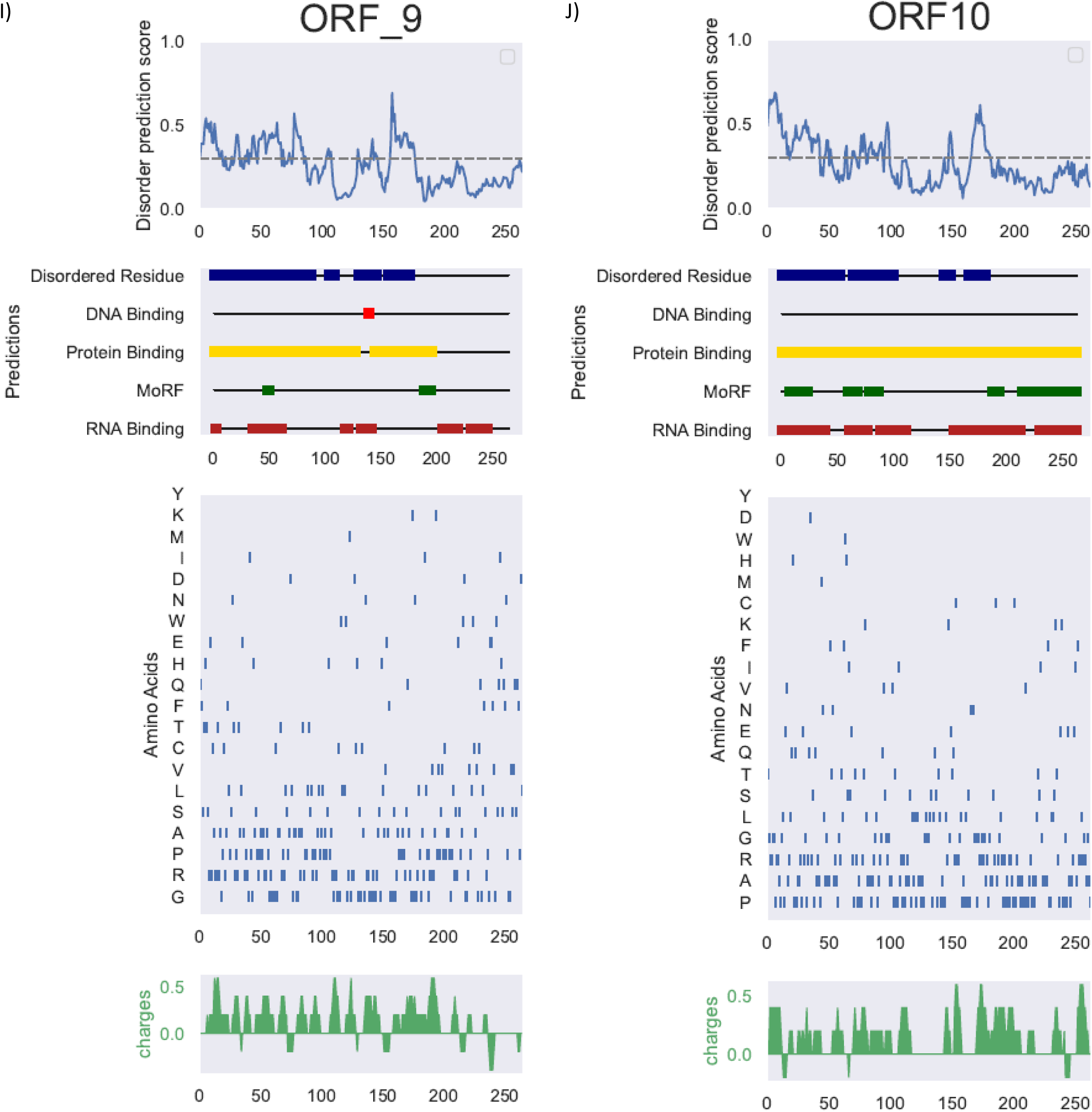
DEPICTER2 results for all 10 eORF-encoded proteins. DEPICTER2 values are presented for all 10 ORFs tested. All results include disorder prediction scores across the amino acid sequence, regions predicted to be disordered residues, MoRF (disorder-to-order transition residues), DNA-binding, RNA-binding, and protein-binding are highlighted in boxes. Additional amino acid frequently detected in the intrinsically disordered regions and charge plots are included for each eORF protein.

**Supplementary Figure 4:**
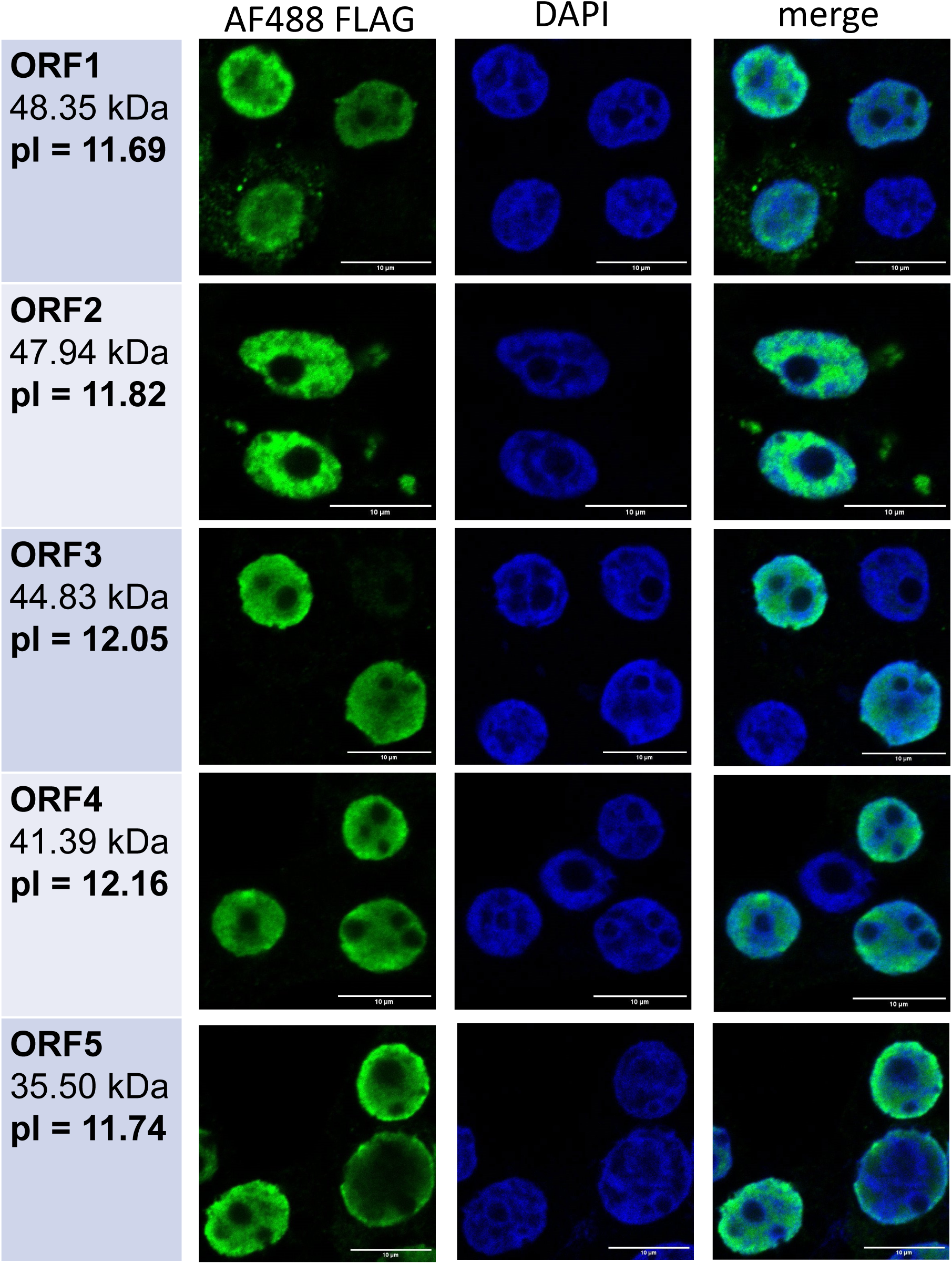
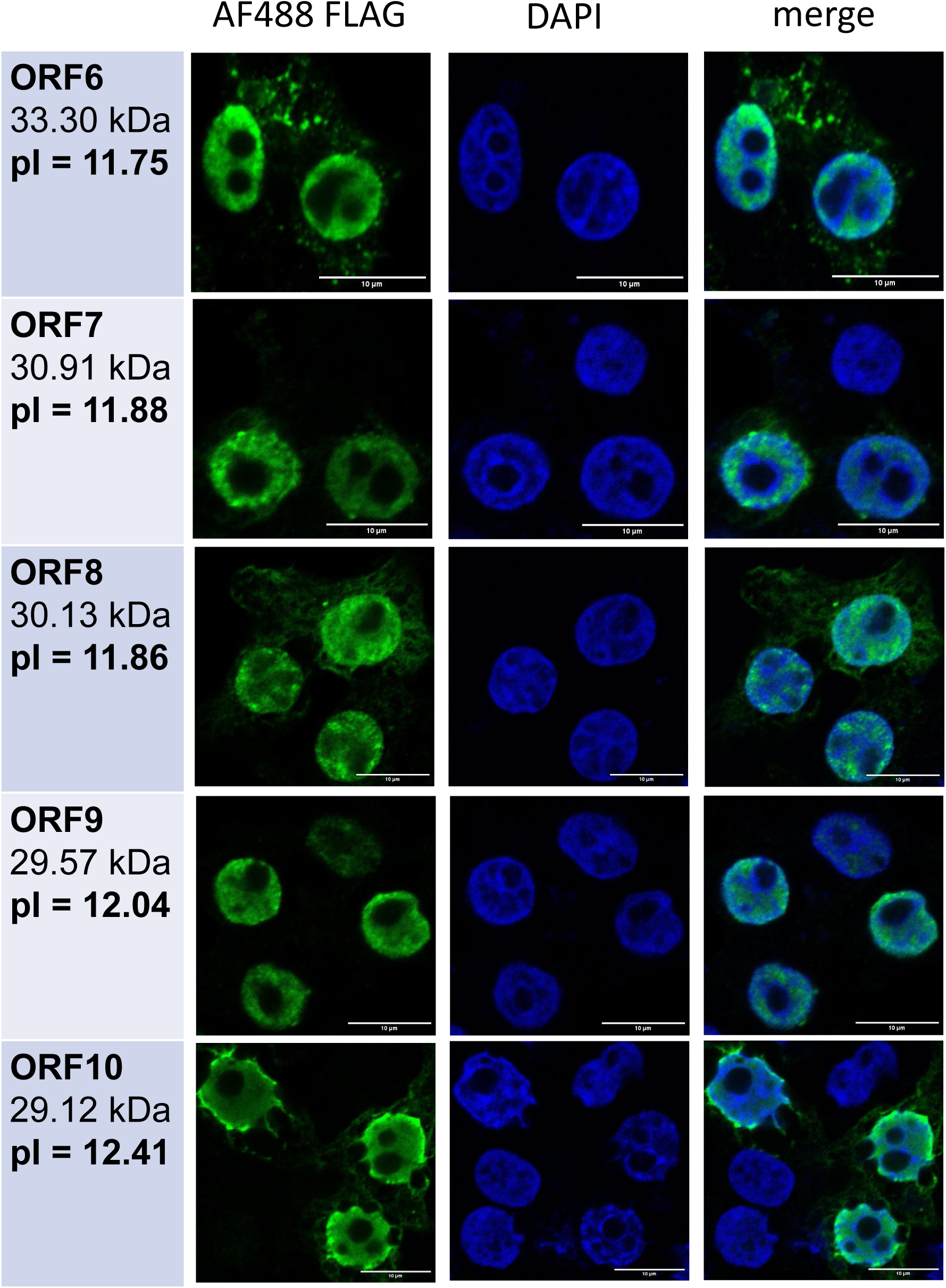
Subcellular localization of eRNA-encoded proteins. Immunofluorescent images of HEK293T cells transfected with plasmids expressing all 10 FLAG-tagged eORF protein constructs. FLAG signal (AF488, green), DAPI (blue) and merged images show localization of four eRNA-encoded proteins. Images from Figure 3A included, and all others used same transfection, fixation, staining, and imaging methods.

**Supplementary Figure 5:**
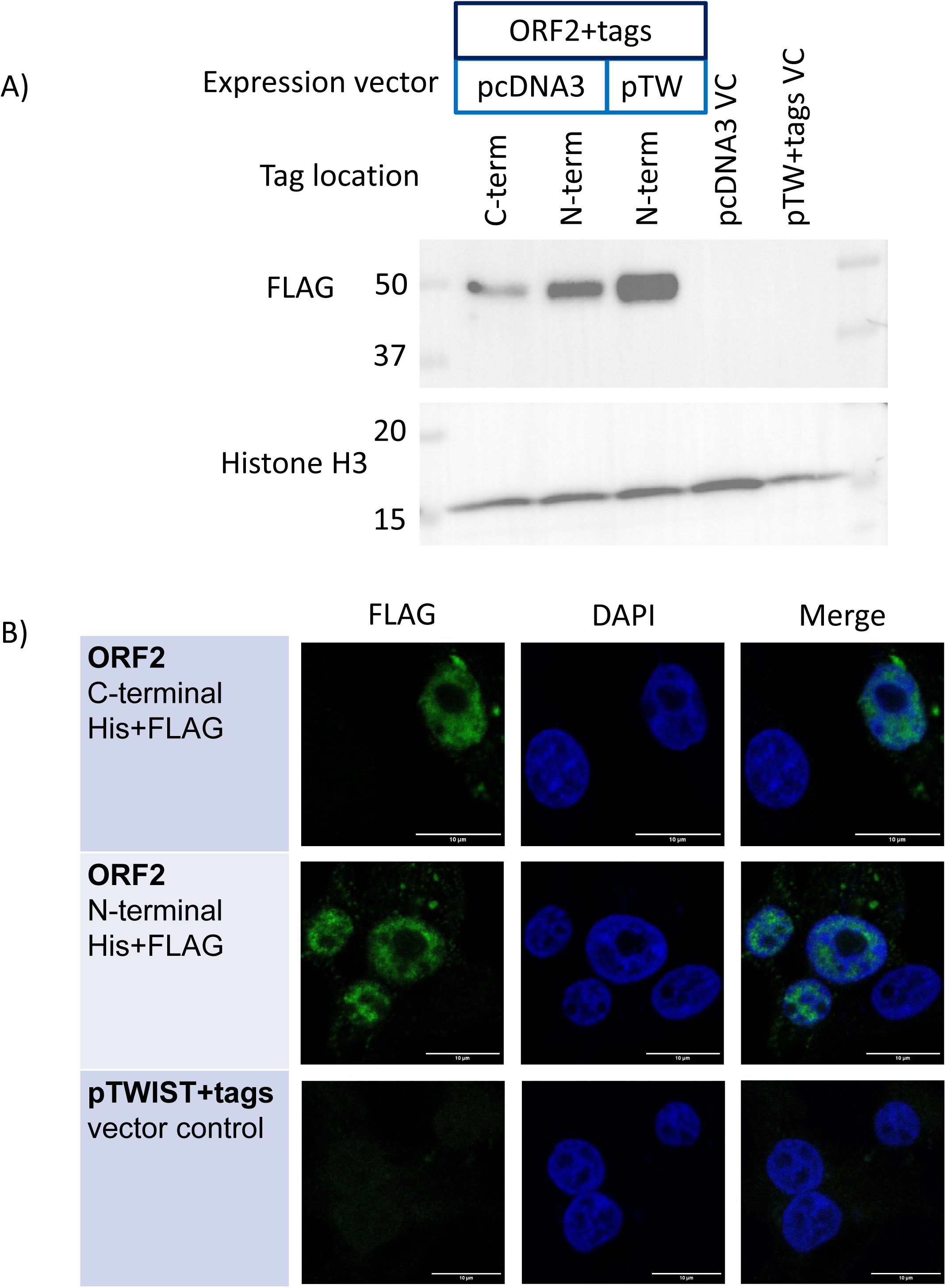
Expression and subcellular localization of C-terminal tagged ORF2 protein cloned from poly-A RNA. Cloned C-terminal tagged ORF2 was expressed in HEK293T cells and compared with its synthesized N-terminal tagged counterpart. All samples were analyzed 48 hours post-transfection. **A** Western blot comparing expression of C-terminally tagged ORF2 versus N-terminally tagged ORF2 in both the pcDNA3 and pTWIST-CMV vectors, with empty pcDNA3 and pTWIST-CMV expressing only the tag sequences as vector controls. Histone H3 was used as a loading control. **B** Immunofluorescent images of cells transfected with pcDNA3 plasmids expressing the C-terminal and N-terminal tagged ORF2 constructs and pTWIST-CMV tag-expressing vector control. FLAG signal (AF488, green), DAPI (blue) and merged images are shown.

**Supplementary Figure 6:**
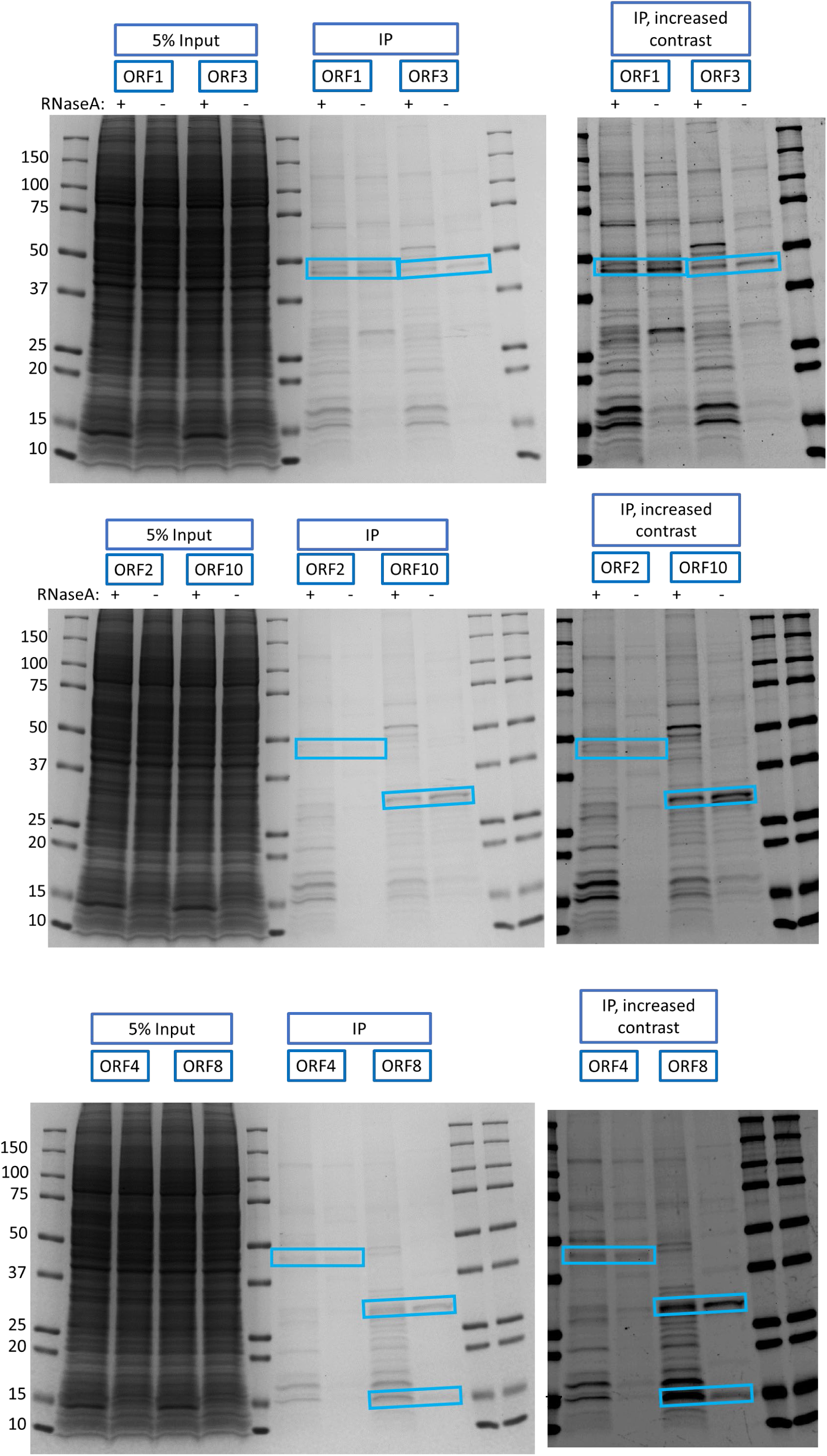
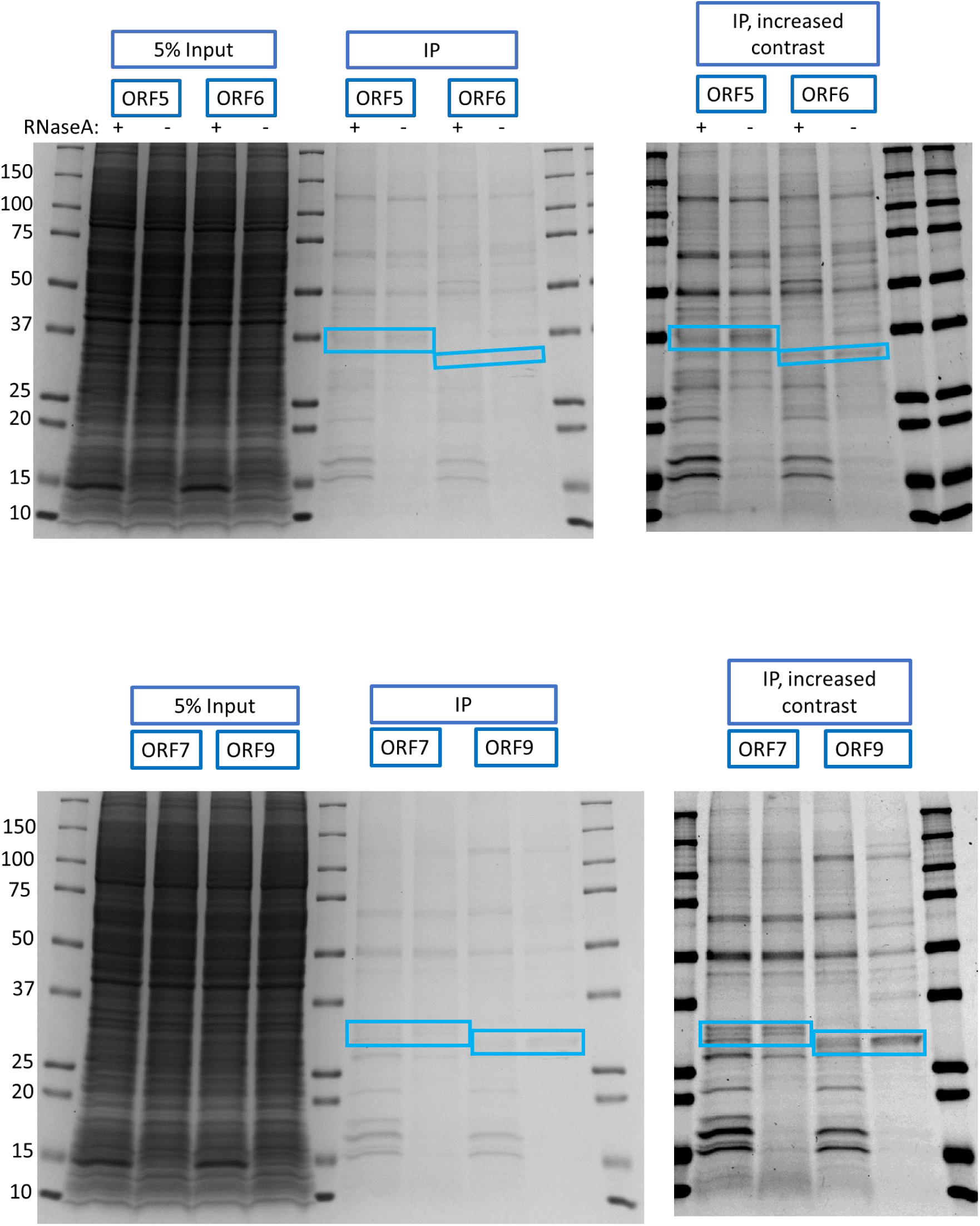
SDS-PAGE and Coomassie staining of coimmunoprecipitated eORFs proteins in RNase-treated and untreated samples. Coomassie-stained SDS-PAGE gels for all 10 tagged eORF proteins isolated by two-stage IP under RNase-treated and RNase-free conditions. Input and IP elution samples are included for all conditions. Additionally, IP lanes are shown with contrast increased using FIJI ImageJ to show band pattern detail. Bands corresponding to FLAG signal confirmed by previous Western Blot experiments are marked with blue.

**Supplementary Figure 7:**
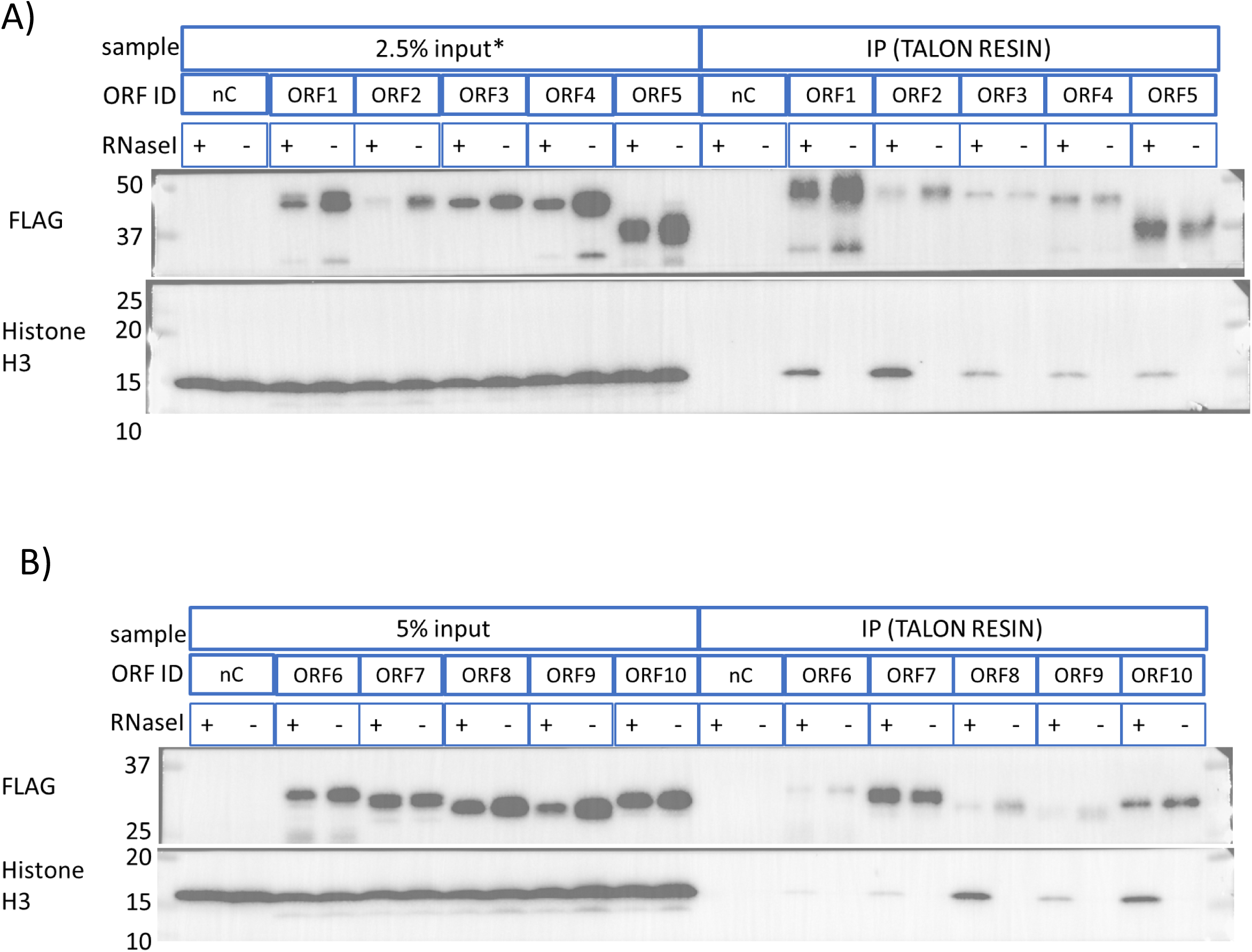
Western blotting confirms presence of histone H3 in eORF protein IP samples. SDS-PAE and Western blotting was carried out with the same samples used in Supplementary Figure 5. Input and IP elution samples are included for each sample, plus an un-transfected negative control (nC). Blots were stained for FLAG tag and Histone H3.

**Supplementary Figure 8:**
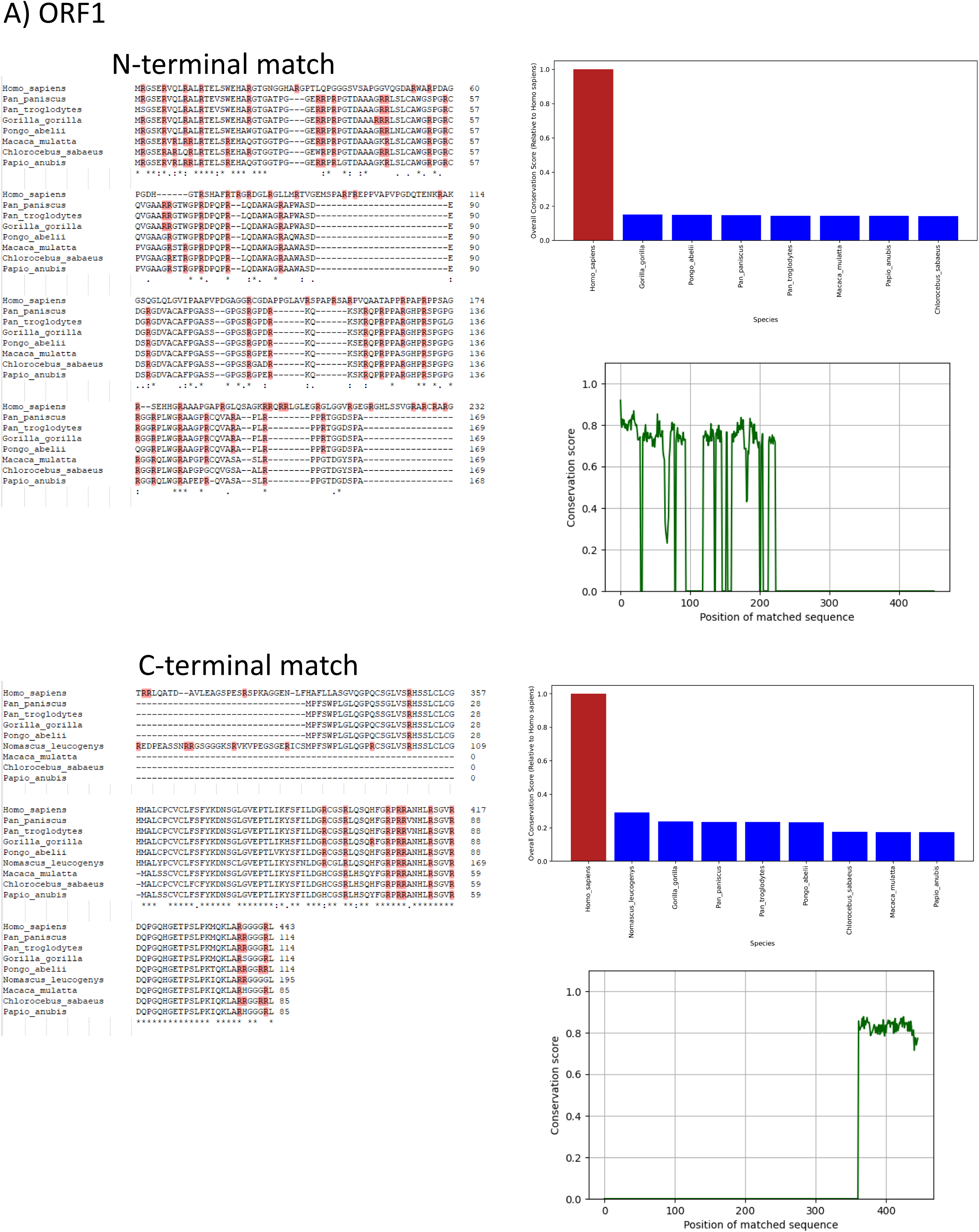
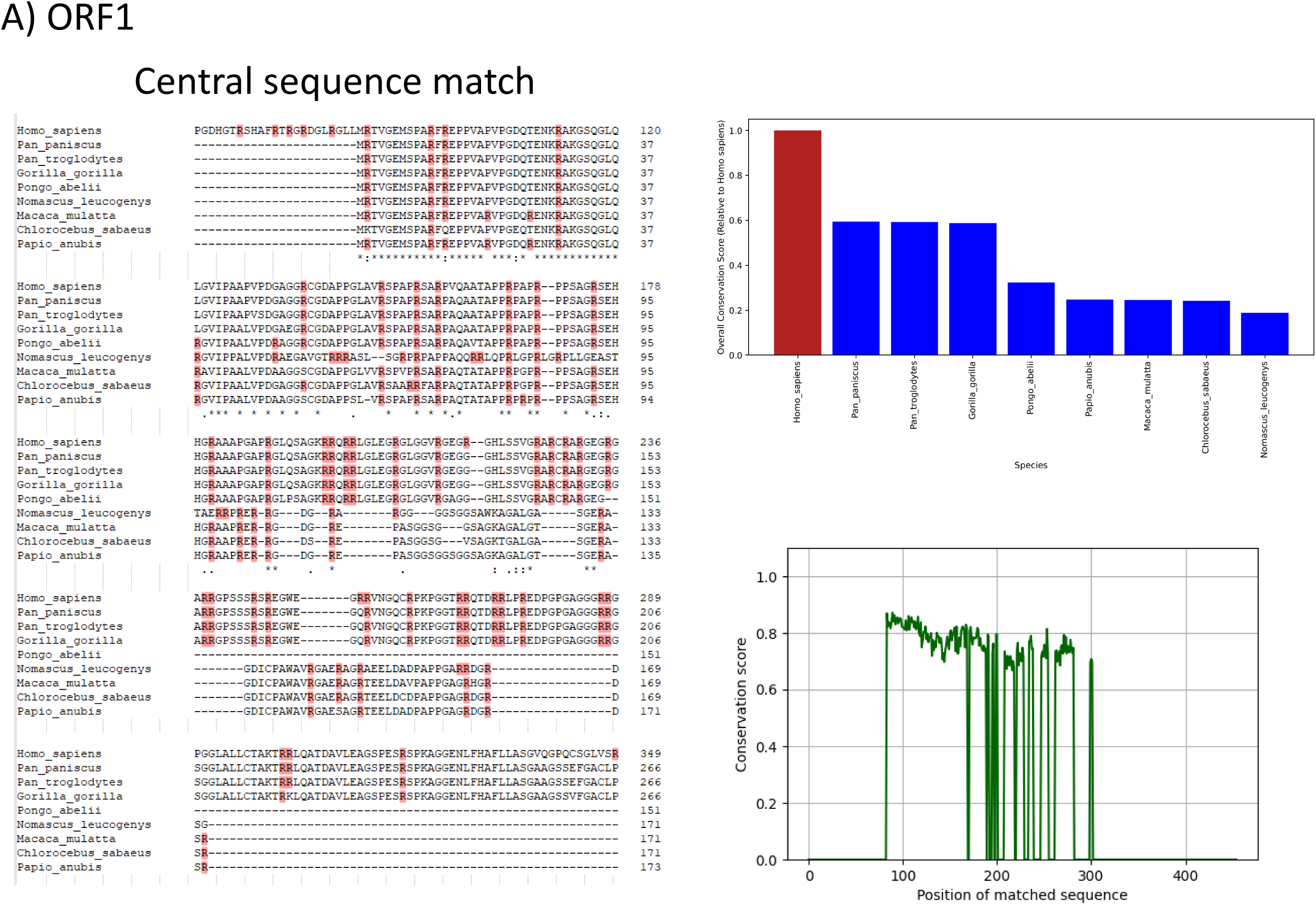
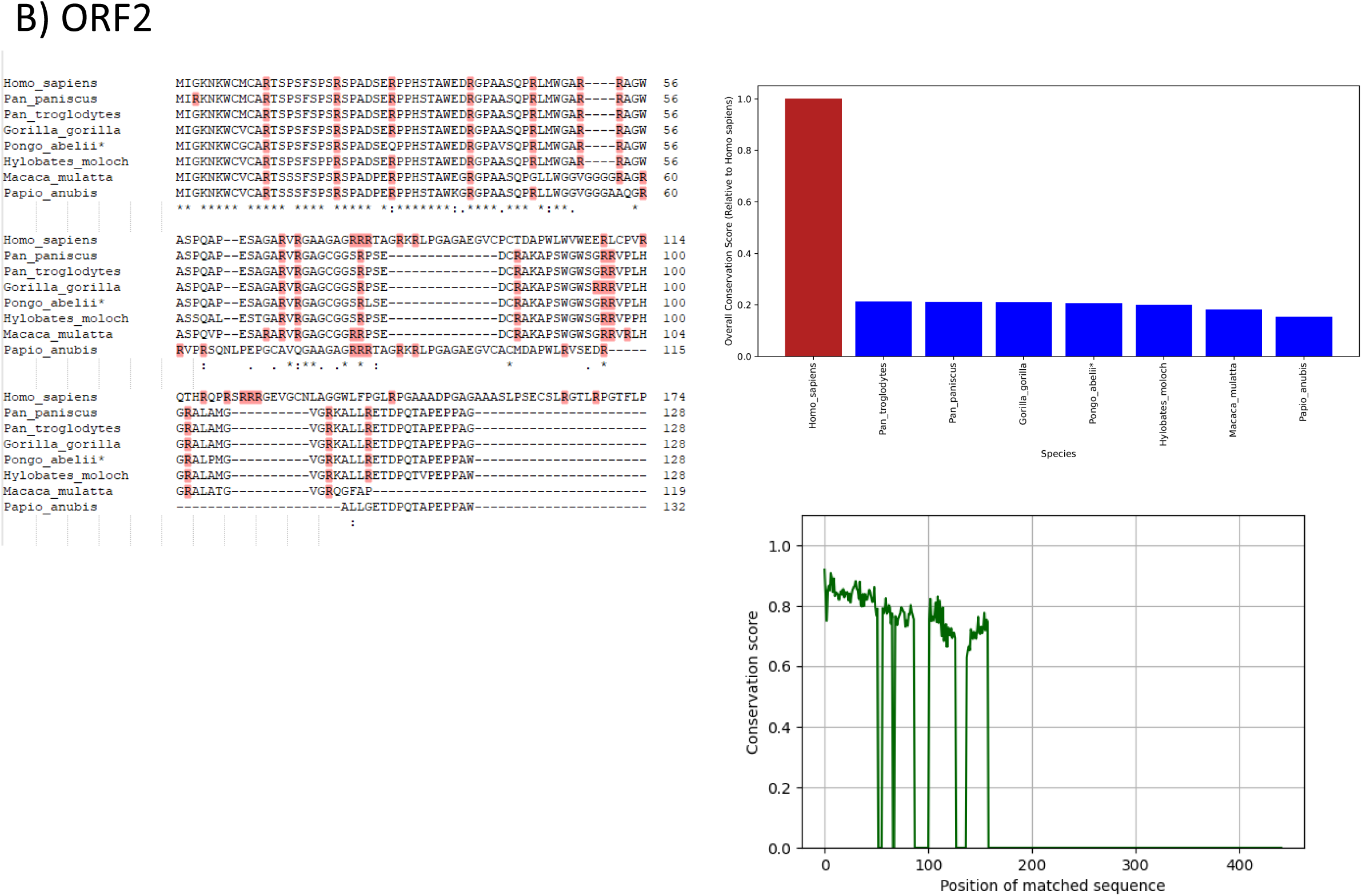
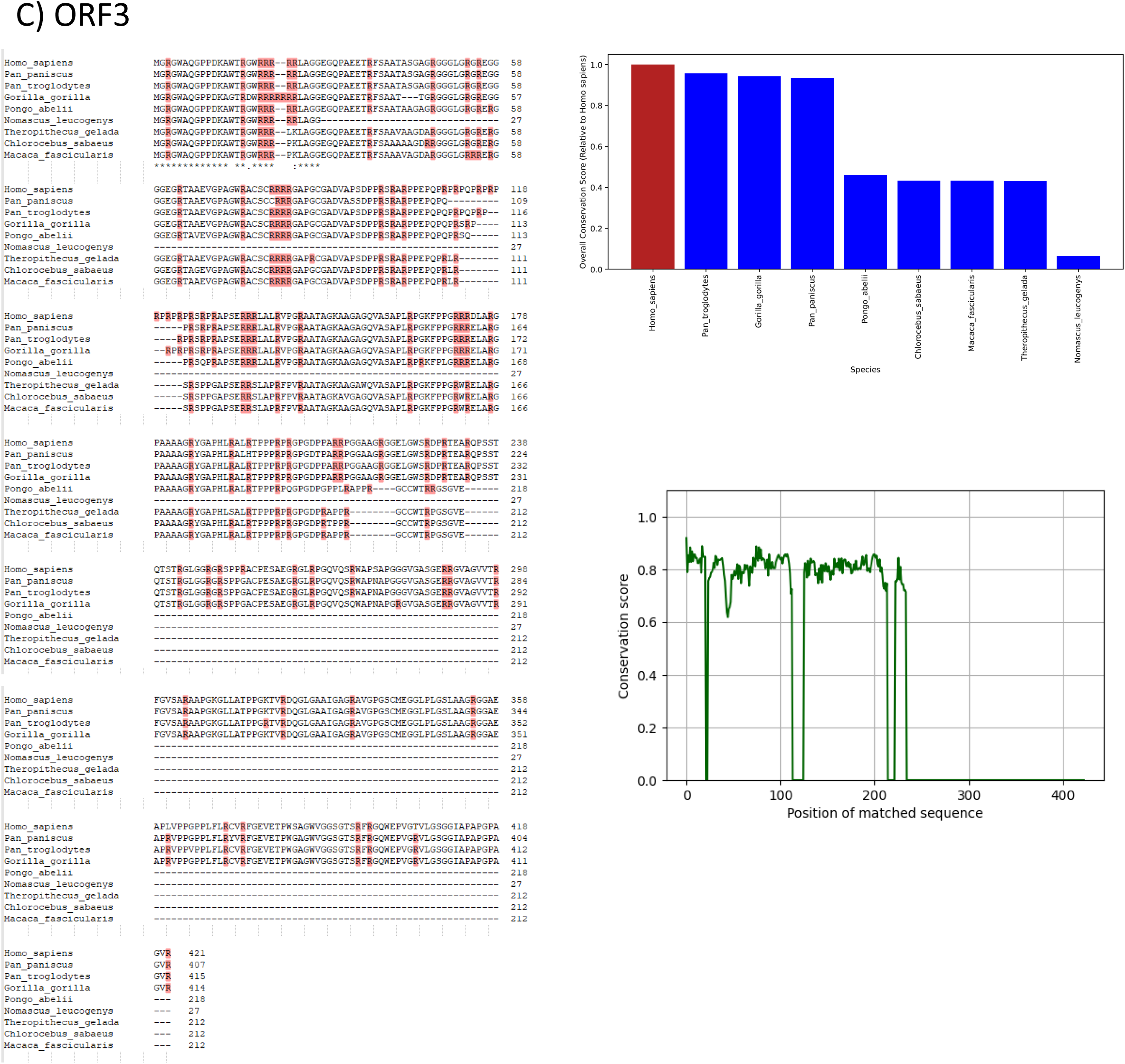
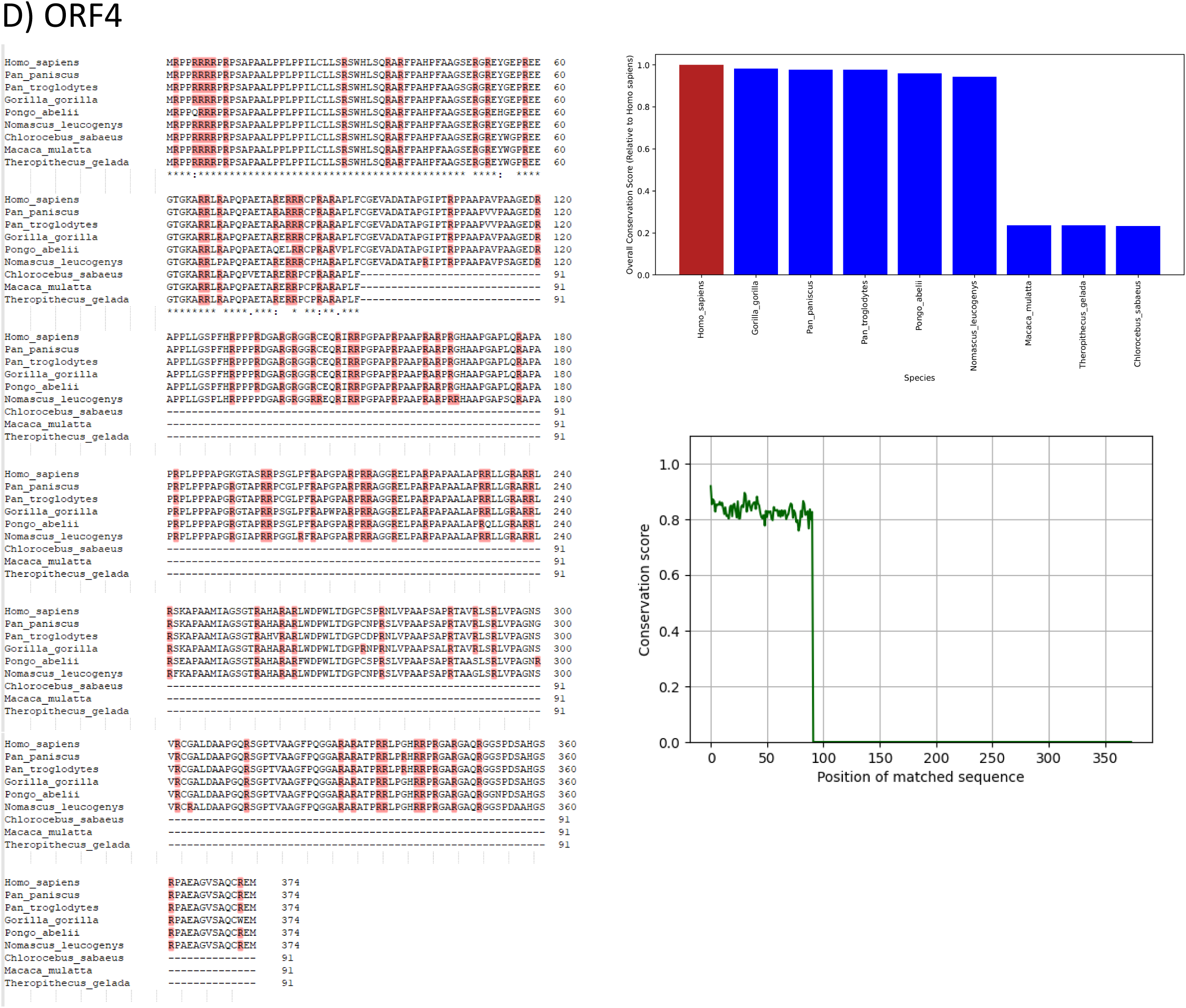
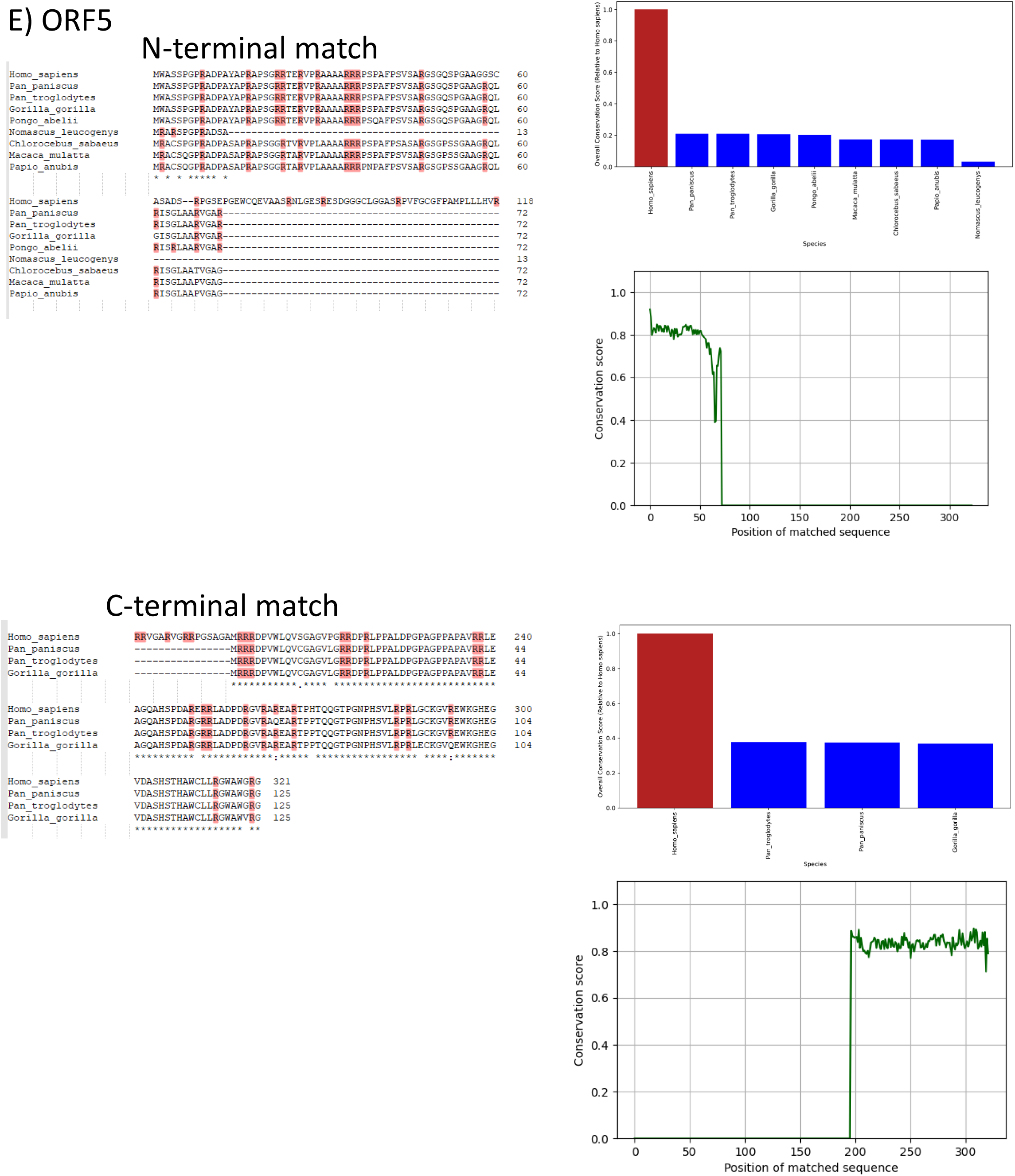
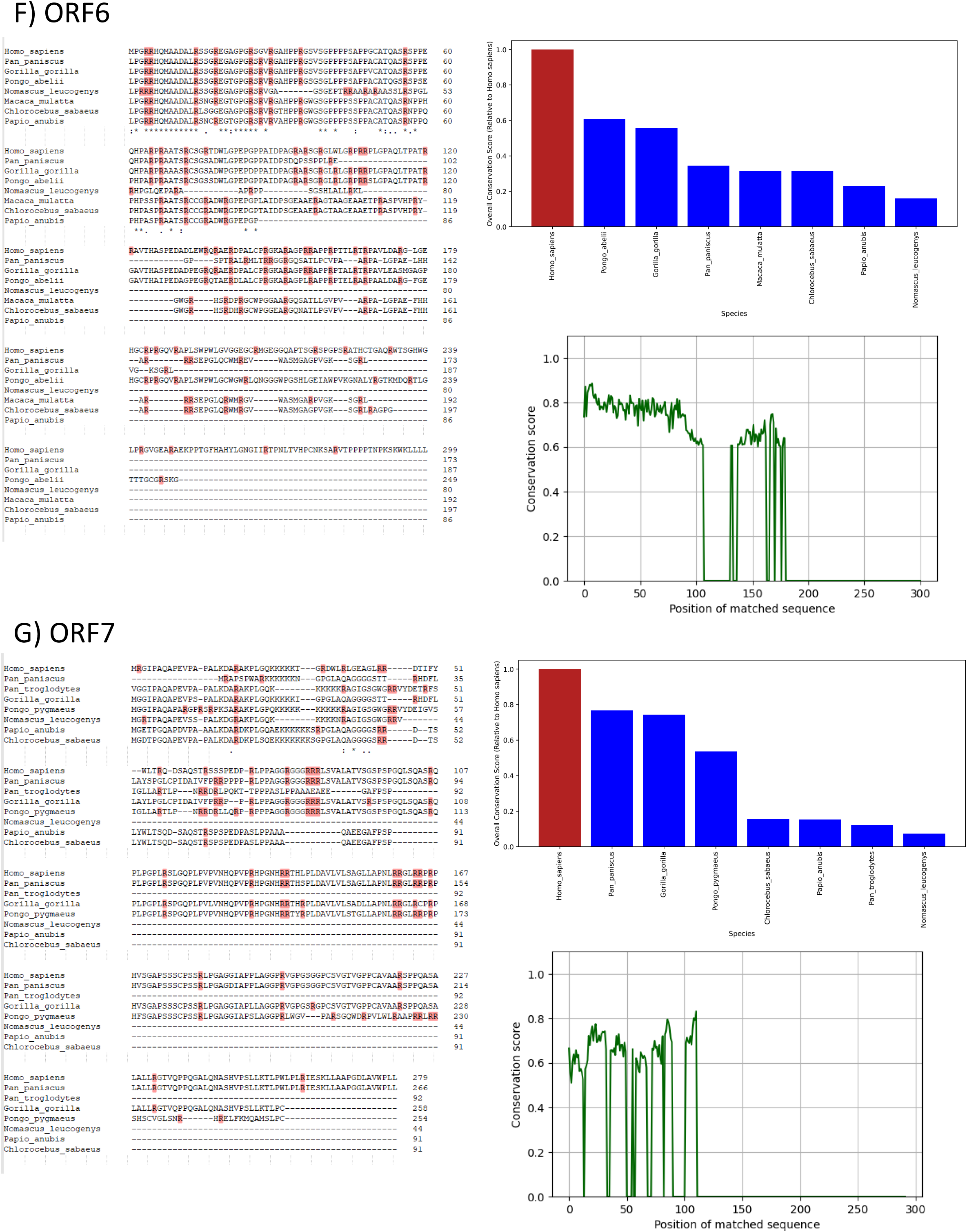
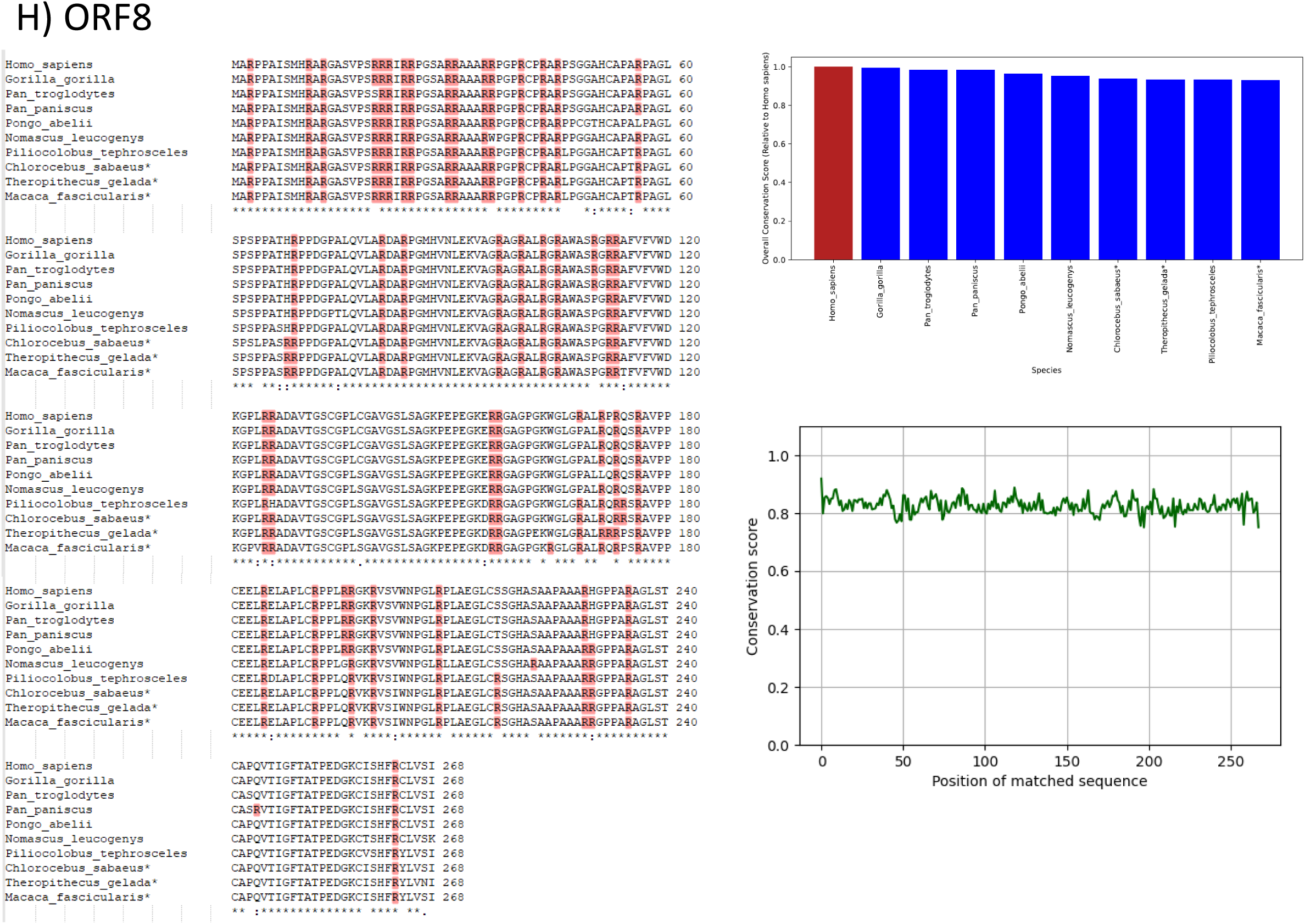
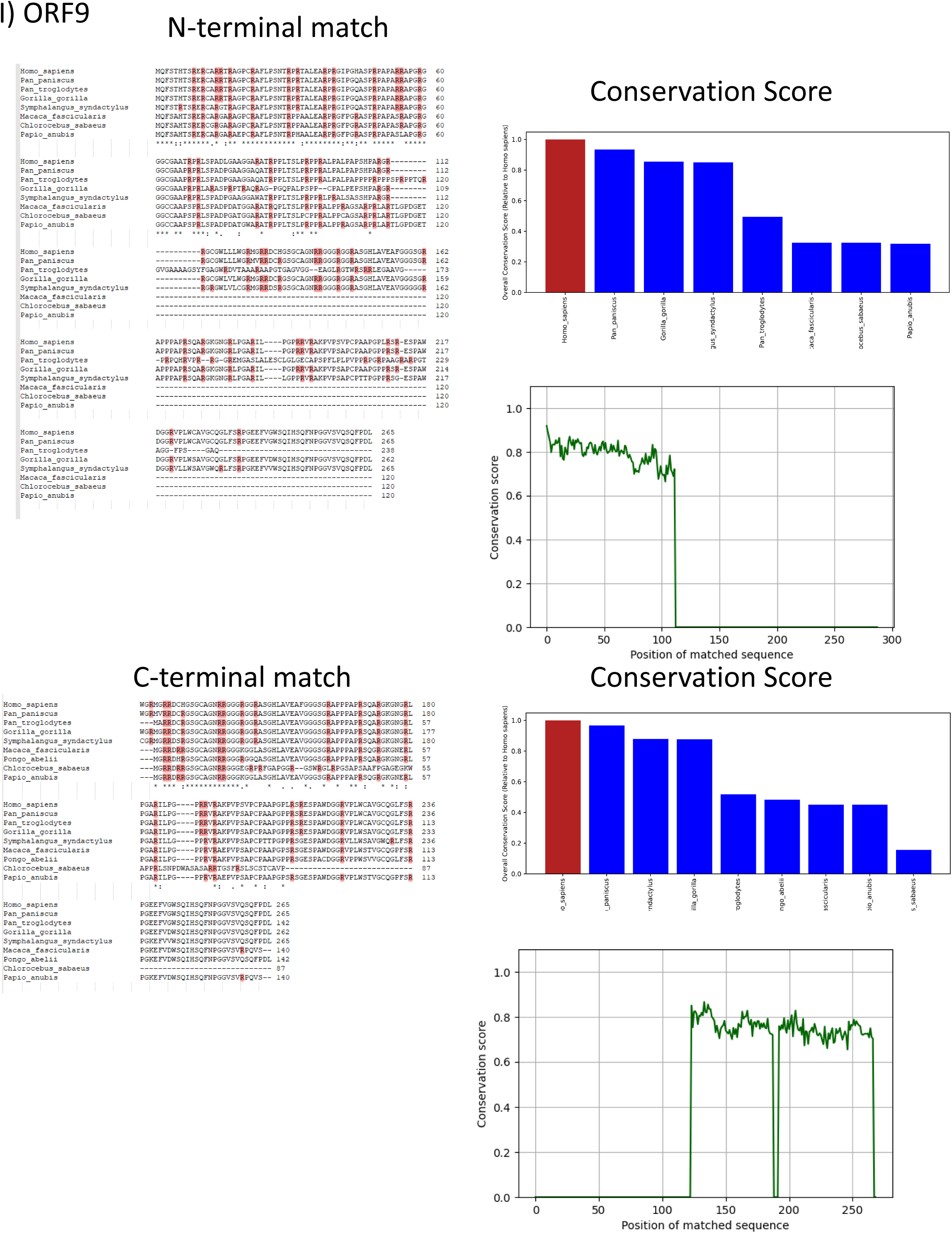
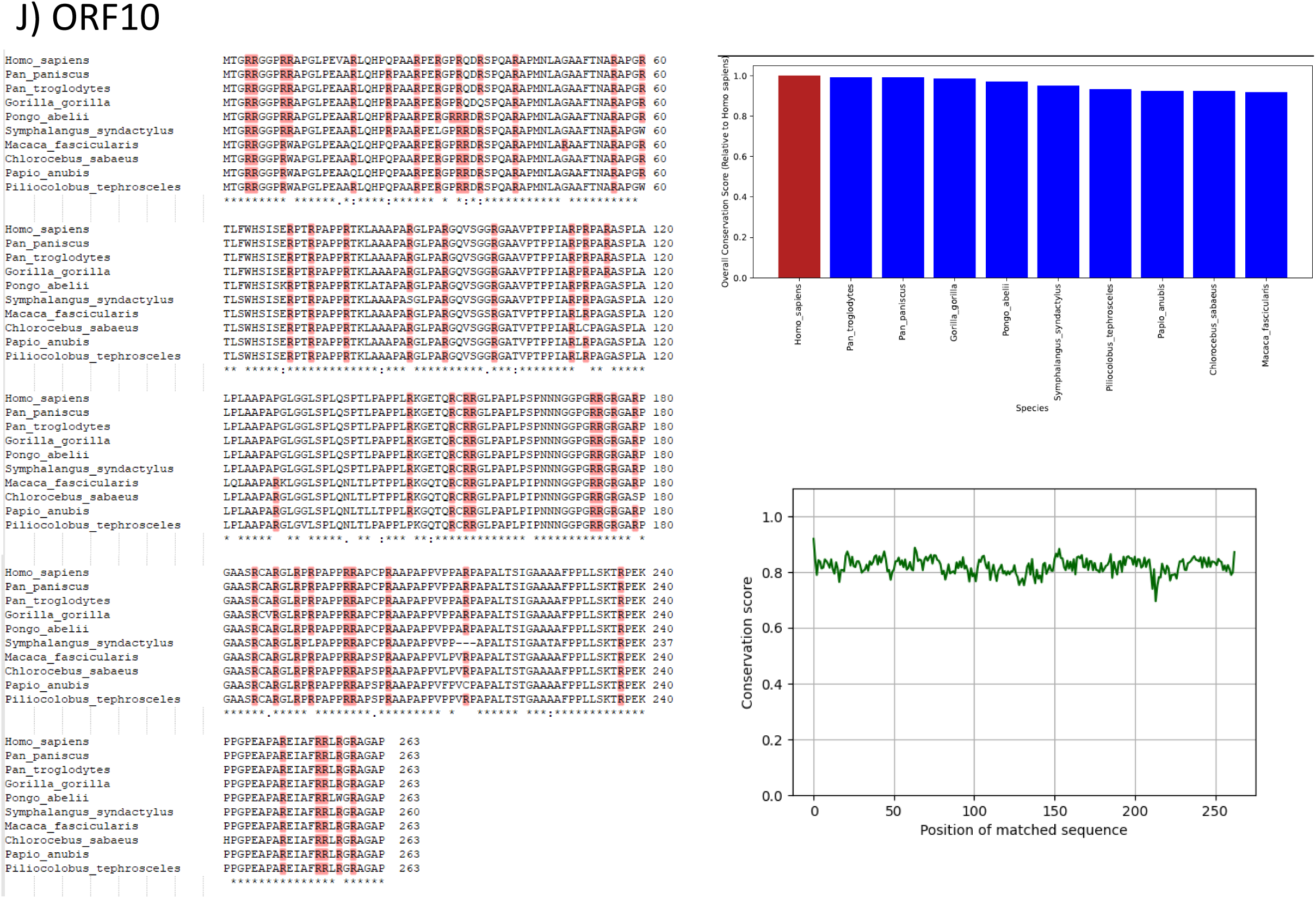
eRNA-encoded proteins share predicted sequences with their ape and monkey homologs. All eORF sequences with their homologs in apes and old-world monkeys are presented, same as in Figure 5A-D. Homologous sequences were identified using NCBI BLASTN and their protein sequences were determined using Expasy Translate. Sequences were aligned using Clustal Omega. Arginine residues are highlighted. Bar plots represent conservation scores for whole sequence, and line plots show per-residue Jensen-Shannon divergence score, calculated in comparison to the full human aa sequence unless otherwise noted. **A** ORF1 non-human homologs contain three ORFs matching the N-terminal, C-terminal, and central regions of the human ORF1 protein. Alignments and conservation plots are presented for all three individually vs the full human sequence. **B** ORF2 non-human homologs only cover the N-terminal region, alignments are presented only for that matching region, conservation plots are shown vs the full human sequence. **C** ORF3 includes homologs matching the full human ORF length. **D** ORF4 includes homologs spanning the full human ORF. **E** ORF5 includes homologs for the N-terminal region across multiple species and C-terminal homologs in homininae only. **F** ORF6 alignments and conservation scores are presented against the full human ORF sequence. **G** ORF7 alignments and conservation scores are presented against the full human ORF sequence. **H** ORF8 homologs shown span the entire human ORF length. **I** ORF9 includes both homologs spanning the entire human ORF and homologs for N- and C-terminal regions, varying by species. **J** ORF10 homologs shown span the entire human ORF length.

**Supplementary Figure 9:**
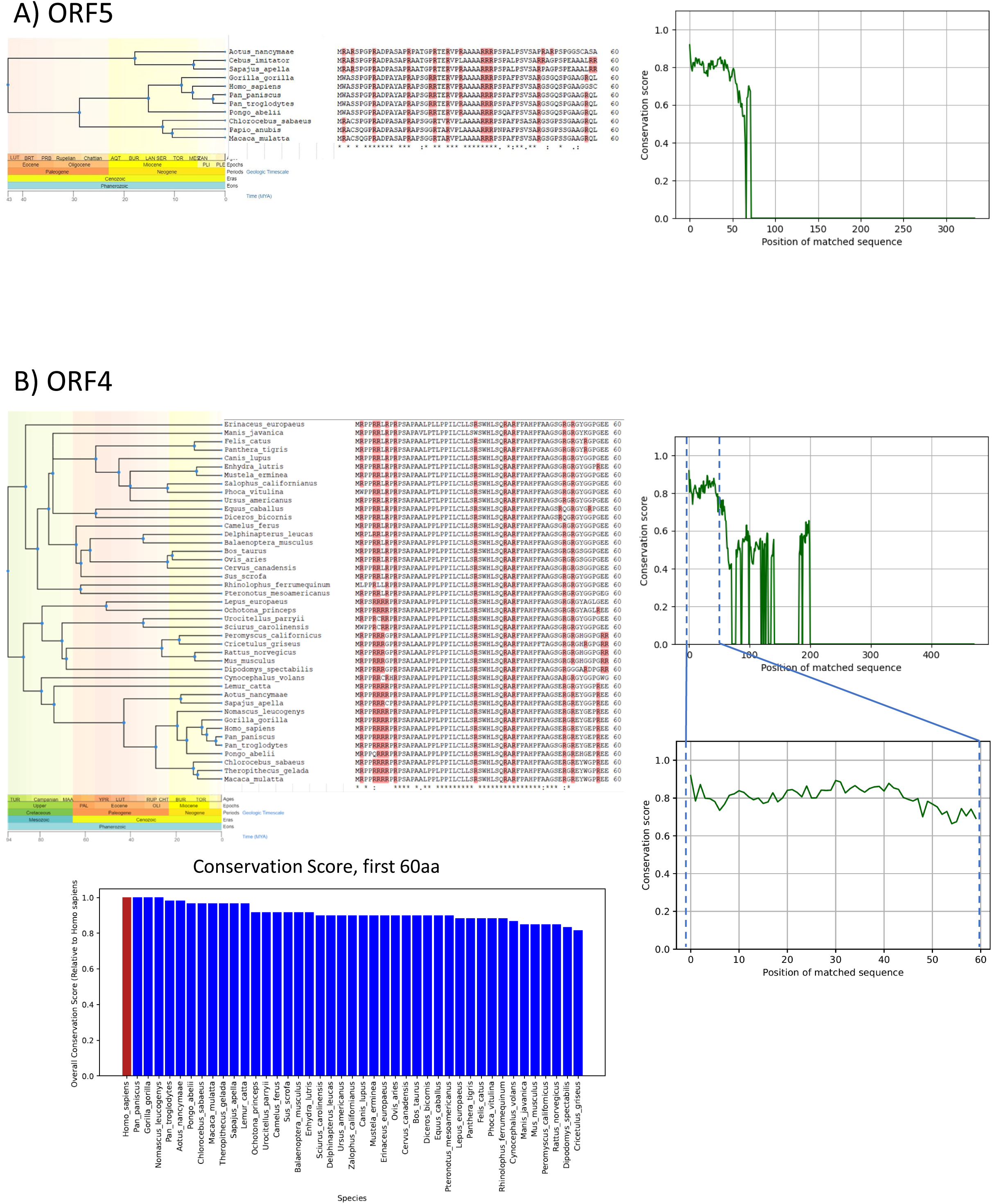
eRNA-encoded proteins display limited amino-acid sequence homology with a wider range of mammal species. ORF4 and ORF5 sequences for a wider range of mammal species were obtained and aligned by the same method as Supp. Fig. 8, with alignments presented limited to the first ∼60 aa showing strong homology between species. Arginine residues are marked. **A** ORF5 alignments and conservation scores are shown for primate species including apes, old-world monkeys, and new-world monkeys. **B** ORF4 alignments and per-residue Jensen-Shannon divergence scores are shown. Additionally, conservation scores for the first 60aa represented in the alignment are shown separately as noted.

